# Genome-Wide Association Study in a Rat Model of Temperament Identifies Multiple Loci for Exploratory Locomotion and Anxiety-Like Traits

**DOI:** 10.1101/2022.07.12.499605

**Authors:** Apurva S. Chitre, Elaine K. Hebda-Bauer, Peter Blandino, Hannah Bimschleger, Khai-Minh Nguyen, Pamela Maras, Fei Li, A. Bilge Ozel, Oksana Polysskaya, Riyan Cheng, Shelly B. Flagel, Stanley J. Watson, Jun Li, Huda Akil, Abraham A Palmer

## Abstract

Common genetic factors likely contribute to multiple psychiatric diseases including mood and substance use disorders. Certain stable, heritable traits reflecting temperament, termed externalizing or internalizing, play a large role in modulating vulnerability to these disorders. To model these heritable tendencies, we selectively bred rats for high and low exploration in a novel environment (bred High Responders (bHR) vs. Low Responders (bLR)). To identify genes underlying the response to selection, we phenotyped and genotyped 558 rats from an F_2_ cross between bHR and bLR. Several behavioral traits show high heritability, including the selection trait: exploratory locomotion (EL) in a novel environment. There were significant phenotypic and genetic correlations between tests that capture facets of EL and anxiety. There were also correlations with Pavlovian conditioned approach (PavCA) behavior despite the lower heritability of that trait.

Ten significant and conditionally independent loci for six behavioral traits were identified. Five of the six traits reflect different facets of EL that were captured by three behavioral tests. Distance traveled measures from the open field and the elevated plus maze map onto different loci, thus may represent different aspects of novelty-induced locomotor activity. The sixth behavioral trait, number of fecal boli, is the only anxiety-related trait mapping to a significant locus on chromosome 18 within which the *Pik3c3* gene is located. There were no significant loci for PavCA. We identified a missense variant in the *Plekhf1* gene on the chromosome 1:95 Mb QTL and *Fancf* and *Gas2* as potential candidate genes that may drive the chromosome 1:107 Mb QTL for EL traits. The identification of a locomotor activity-related QTL on chromosome 7 encompassing the *Pkhd1l1* and *Trhr* genes is consistent with our previous finding of these genes being differentially expressed in the hippocampus of bHR vs. bLR rats.

The strong heritability coupled with identification of several loci associated with exploratory locomotion and emotionality provide compelling support for this selectively bred rat model in discovering relatively large effect causal variants tied to elements of internalizing and externalizing behaviors inherent to psychiatric and substance use disorders.

## Introduction

Psychiatric epidemiology studies suggest that common genetic factors underlie multiple stress-related psychiatric diseases including mood and substance use disorders. These complex disorders have been placed into two broad categories termed “internalizing disorders” or “externalizing disorders” (Cerdá et al., 2010; Kendler, 1992), with certain traits or stable aspects of temperament being central to the vulnerability to these two classes of psychopathology. Specifically, two personality traits measured with formal psychological instruments have emerged as key predictors: *neuroticism* is a strong predictor of internalizing disorders, such as depression, anxiety and other mood disorders (Bienvenu et al., 2001; Clark et al., 1994; Jardine et al., 1984; Khan et al., 2005), and *high sensation seeking* is a predictor of externalizing disorders, including conduct disorder, the propensity for risky behaviors and drug use (Cloninger, 1987; Conway et al., 2003; Etter et al., 2003; Zuckerman & Cloninger, 1996; Zuckerman & Kuhlman, 2000). These temperamental vulnerabilities emerge very early in development. (Kagan & Snidman, 1999). For instance, toddlers with high levels of behavioral inhibition show increased risk for internalizing disorders (Biederman et al., 2001; Caspi, 1996; Hayward et al., 1998; Muris et al., 2001; Schwartz et al., 1999), whereas toddlers who are impulsive are at greater risk of developing externalizing disorders (Eigsti et al., 2006). Notably, extremes in both temperaments can interact with environmental triggers and lead to substance use disorders — thus substance use and misuse can be associated with externalizing tendencies that lead to greater experimentation with drugs, or internalizing vulnerabilities where drug use can be viewed as a form of self-medication. Uncovering the genetic mechanisms associated with these temperamental tendencies represents an entry point for elucidating the biological underpinning of these brain disorders and using that knowledge to better treat and prevent them.

Major depressive disorder is heterogenous and moderately heritable (Akil et al., 2018). A meta-analysis of genome wide association studies (GWAS) which comprised a large number of individuals (over 800,000), estimated the genome-wide SNP-based heritability on the liability scale as 0.089 (s.e. = 0.003) and identified 102 variants associated with this illness (Howard et al., 2019). The existence of a genetic component to addiction vulnerability has been well known for over two decades (Merikangas et al., 1998) and addiction to various substances is moderately to highly heritable, as observed in large twin studies (Goldman et al., 2005) and more recently in large GWAS studies (Gelernter & Polimanti, 2021). Despite this progress, identification of causal allelic variants and genes has been difficult due to the complex nature of substance use disorders and the comorbidity among substance use and other psychiatric disorders. GWAS are beginning to address these knowledge gaps. Until recently, the only risk genes that were well established for substance use disorders acted pharmacogenetically: alcohol-metabolizing enzyme genes for alcohol traits, nicotinic receptors for nicotine traits, and μ-opioid receptor for opioid traits (Gelernter & Polimanti, 2021). Recent large GWAS studies have identified loci that are associated with the use and abuse of substances including alcohol, nicotine, and more recently cannabis and opiates (reviewed in (Gelernter & Polimanti, 2021)). A key insight that emerges from these recent addiction-related GWASs is the importance of phenotypic traits related to quantity and/or frequency measures of substance use, versus symptoms related to physical dependence, as best established with respect to alcohol.

In parallel, GWAS of various temperamental traits including neuroticism (reviewed in (Sanchez-Roige et al., 2018)), impulsivity (Sanchez-Roige et al., 2019), and externalizing behaviors (Karlsson Linnér et al., 2021) have established robust genetic correlations between temperament, mood disorders, and substance abuse. Neuroticism has been the best studied personality trait with several loci identified via GWAS and strong positive genetic correlations with major depressive and anxiety disorders (Sanchez-Roige et al., 2018). Recently, two genetically distinguishable subclusters of neuroticism, ‘depressed affect’ and ‘worry’, have been identified, suggesting distinct causal mechanisms for subtypes of individuals (23andMe Research Team et al., 2018). Impulsivity has been increasingly recognized as a heterogeneous phenotype, with both overlapping and unique genetic substrates for individual differences in impulsive personality traits (Sanchez-Roige et al., 2019). Notably, sensation-seeking, the preference for highly stimulating experiences, is genetically distinct from the other impulsivity traits. There is also genetic overlap between various measures of impulsivity and several psychiatric conditions, including substance use disorders. Several GWAS of the group of correlated traits called externalizing have recently been examined using a multivariate approach revealing a polygenic score accounting for 10% of the variance in externalizing, one of the largest effect sizes of any polygenic score in psychiatric and behavioral genetics (Karlsson Linnér et al., 2021). These recent large GWAS, and associated meta-analyses, of temperamental traits are beginning to reveal the genetic underpinnings of individual differences in vulnerability to psychiatric and substance use disorders.

One advantage of the focus on temperament is that, compared to psychiatric diagnoses, temperamental traits can be better modeled in rodents. These models can then be used to parse the genetic, developmental, and environmental factors that contribute to behavioral phenotypes and to directly investigate the associated neural mechanisms. To this end, we have been selectively breeding Sprague-Dawley rats for the sensation-seeking trait as manifested by their propensity to explore a novel, mildly stressful environment (Stead et al., 2006).

Novelty-induced exploratory locomotion (EL) can be indexed using a constellation of measures, including total distance traveled in the open field (OF) and the elevated plus maze (EPM) as well as lateral and rearing locomotor scores in the OF. Distance traveled in the OF has been previously associated with differential vulnerability to drug self-administration in outbred rats (Piazza et al., 1989). Although selected exclusively based on EL, we tested a number of other spontaneous behaviors as well as stress- and addiction-related behaviors to see if they were genetically correlated with the selection trait. As they diverged, our two lines came to represent the polar extremes of emotionality exhibited by the outbred Sprague-Dawley rats from which they are derived (Flagel et al., 2014; Stead et al., 2006; Turner et al., 2017). The bred high responders (bHRs) exhibit externalizing behaviors of sensation seeking and impulsivity, low anxiety-like behavior, greater propensity to psychostimulant sensitization, and lower thresholds for drug- and cue-induced relapse when compared to both outbred Sprague-Dawley rats and the bred low responders (bLRs) (Aydin et al., 2021; Flagel et al., 2016). In turn, bLRs exhibit internalizing anxiety- and depressive-like behavior and are more responsive to psychosocial stress, which triggers drug-seeking behavior at a level comparable to the bHRs (Clinton et al., 2014). bHRs and bLRs also respond very differently to environmental stimuli associated with reward during a Pavlovian conditioned approach (PavCA) task; a cue associated with reward becomes a reinforcer by itself for the bHRs (“sign-tracking”), while a cue serves only as a predictor of reward for bLRs (“goal-tracking”) (Flagel et al., 2011). The highly divergent phenotypes of the bHR and bLR rats reveal how these lines exemplify two distinct and clinically relevant paths to substance abuse: sensation seeking and high reactivity to psychosocial stress. These behavioral phenotypes of the bred lines are robust and stable beginning early in development (Clinton et al., 2014; Turner et al., 2017), similar to temperament in humans (Mervielde et al., 2005).

In the present study, we genotyped 20 F_0_ and 558 F_2_ rats that were produced by intercrossing bHR and bLR rats from the 37th generation of selection. Because bHR and bLR rats are outbred (Stead et al., 2006; Zhou et al., 2019), the genetic analysis of this cross was more complex than an F_2_ cross between two inbred strains, and therefore it required much denser marker coverage. We performed a GWAS for several behaviors that model various aspects of these phenotypes: sensation seeking, anxiety behaviors, and PavCA behaviors. Our findings shed light on genetic differences associated with specific behavioral characteristics of these two lines.

## Methods

### Phenotyping

Rats selectively bred for a high or low propensity to explore a novel environment (bHRs and bLRs) at two months of age were established at the Michigan Neuroscience Institute, University of Michigan (Stead et al, 2006). The two lines of rats exhibit predictable differences in exploratory and anxiety-like behaviors (Flagel et al., 2014). After 37 generations of selective breeding, an initial set (“F_0_”) of bHRs deriving from 12 distinct families were bred with bLRs from 12 distinct families to create 12 intercross families. The offspring of this intercross (“F_1_”) were then bred with each other to help the re-emergence/divergence of the phenotypes in the F_2_ generation. To amplify divergence, F_1_ rats that had a lower response to novelty were bred together, and F_1_ rats that had a higher response to novelty were bred together. The resulting 48 F_2_ litters generated a total of 538 male and female rats, of which 216 were tested as adolescents (about 1 month old) in the open field, since our typical locomotor response to a novel environment test is geared towards adult rats, and sacrificed at this age to collect tissue. The remaining 322 rats grew to young adulthood at which time they were phenotyped and then sacrificed for tissue collection (about 4 months old). The ages, sex, and number of rats phenotyped for each behavioral test are shown in **Table 1**.

**Table 1.** Number of rats phenotyped Number of F_2_ rats phenotyped. Summary of the number of rats phenotyped for each test, as well as ages when phenotypes were collected

| Behavioral Test | Males, N | Females, N | Total N | Age, days |
| --- | --- | --- | --- | --- |
| Locomotor Response to a Novel Environment | 171 | 151 | 322 | 123 +/- 5 |
| Elevated Plus Maze | 171 | 151 | 322 | 123 +/- 5 |
| Pavlovian Conditioned Approach | 106 | 105 | 211 | 123 +/- 4 |
| Open Field | 103 | 113 | 216 | 27 +/- 2 |

All rats were weaned at postnatal day 21, housed 2-3 rats per standard rat cage (140 sq. inches), and provided ad lib water and standard rat chow (5LOD, PicoLab Laboratory Rodent Diet, LabDiet) in a temperature (70-78°F) and humidity (30-50%)-controlled room with a 12/12 light/dark cycle (lights on at 0600, lights off at 1800). All procedures were conducted in accordance with the guidelines outlined in the National Institutes of Health Guide for the Care and Use of Animals and were approved by the Institutional Animal Care and Use Committee at the University of Michigan.

#### Adolescent Behavioral Testing

In adolescents, EL and anxiety-like behaviors were measured using an open field (OF) test. The OF is a 100 cm × 100 cm × 50 cm white Plexiglas box with a matte black floor. All testing was performed between 9:00 am and 4:00 pm under dim light conditions (40 lux). The test began by placing each rat into one corner of the OF and then allowing five minutes for free exploration of the apparatus. During the test, the rat’s behavior was monitored via a video tracking system (Ethovision, Noldus Information Technology). This tracking system was used to sub-divide the floor of the OF into a total of 16 zones: the inner four zones were defined as the anxiogenic region of the apparatus; the time spent in these inner zones was recorded (converted to % time of five min). Additional measures during the test (i.e, total distance traveled in cm, duration of immobility in sec) were also recorded. The number of fecal boli was also counted as an anxiety-related index (Hall, 1938).

### Adult Behavioral Testing

#### Locomotor Response to a Novel Environment

Adult rats were initially tested at approximately two months of age to assess their EL in a novel environment that was the same size as a home cage (23 cm × 43 cm) but located in a different room with novel cues. The number of beam breaks (counts) was measured over 60 minutes for both lateral and rearing movements, deemed the lateral and rearing locomotor scores. The cumulative lateral and rearing locomotor scores over the 60-minute test were combined to generate the total locomotor score.

#### Elevated Plus Maze (EPM)

Spontaneous anxiety-like behavior was evaluated in the EPM, which consists of four black Plexiglass arms (45 cm long, 12 cm wide) elevated 70 cm from the floor, arranged in the shape of a cross. Two opposite arms are enclosed by 45 cm high walls, and the other two opposite arms are open. The test began by placing animals in the intersection of these arms, a 12 × 12 cm square platform which allows access to all four arms. During the five-minute test period, the room is dimly lit (40 lux) and animals’ behavior monitored using a video tracking system (Ethovision, Noldus Information Technology) that records the amount of time spent in the arms (converted to % time of five min), the distance traveled in cm, and the time immobile in sec. As in the OF for the adolescent rats, the number of boli was counted as an additional measure of anxiety.

#### Pavlovian Conditioning

In addition to EL and anxiety-like differences that have emerged in the selective breeding of the bHR and bLR animals, these two lines show stark differences in their PavCA, with the bHRs showing preference for approaching the reward cue (sign-trackers; ST) and the bLRs showing preference for approaching the location of reward delivery (goal-trackers; GT) (Flagel et al., 2011). Thus, a subset of the adult F_2_ rats were later tested for PavCA behavior, as described in (Meyer et al., 2012), to determine the re-emergence of this phenotype. Briefly, rats were exposed to 7 sessions of PavCA training, each of which consisted of 25 pairings of a lever-cue (conditioned stimulus) followed by (non-contingent) delivery of food reward (unconditioned stimulus). Three dependent variables are averaged to generate a PavCA Index: a) probability difference, which is the difference in the probability to approach the lever-cue vs. food cup during cue presentation; b) response bias, which is the difference in the number of lever-cue contacts vs. food cup entries; and c) latency score, which is the difference in latency to approach the lever-cue vs. the food cup. The PavCA Index for the last two days of testing (i.e., days 6 and 7 for the current study) are used to determine if animals primarily display sign-tracking (values >0.5) or goal-tracking (values <−0.5) behavior. However, we also included the intermediate responders for correlational analyses in order to capture the full population.

### Genotyping via lcWGS

High molecular weight (HMW) genomic DNA from 538 F_2_ spleen samples was isolated via the MagAttract HMW DNA Kit (Qiagen) at the University of Michigan and transferred to the University of California San Diego for further processing. Multiplexed sequencing libraries were prepared using the RipTide kit (iGenomX), and then sequenced on a NovaSeq 6000 (Illumina), according to the manufacturer’s instructions. We produced an average of ~2.8 million reads per sample. The reads were aligned to the rat reference genome Rnor_6.0 (rn6) from Rat Genome Sequencing Consortium (GCA_000001895.4 GCF_000001895.5).

Because single nucleotide polymorphisms (SNPs) that segregate between bHR and bLR were not known, we deeply sequenced 20 rats from the F_0_ generation to identify SNPs to genotype in the larger cohort. For these 20 rats, liver tissue was sent to HudsonAlpha Discovery (Discovery Life Sciences) for HMW genomic DNA extraction and long-range and deep sequencing. Following 10X Chromium linked-read sequencing library preparation, samples were sequenced on one NovaSeq 6000 (Illumina) S4 PE150 flowcell. An average of 999.8 million reads per sample were generated, with an average coverage of 47.25X. Variants in the 538 F_2_ rats were called using a two-step procedure. First, we used STITCH v1.6.6 (R. W. Davies et al., 2016), which is designed to impute high quality genotypes from low coverage sequencing data. We set niteration parameter to 1 and used the SNPs from the 20 deeply sequenced F_0_ rats as reference data. After STITCH was run, only SNPs with INFO score > 0.65 and ‘missing’ rate < 0.15 were retained. Next, we used BEAGLE v5.2 (Browning & Browning, 2007), to impute missing genotypes.

Once the genotypes were obtained, we performed several quality control steps to identify potential sample mix-ups due to sample handling errors: (1) we compared genotypes to the earlier microarray-derived genotypes (Zhou et al., 2019) that were available for 319 of the rats in the present study, (2) we assessed the sex, as determined by the fraction of reads mapping to the X and Y chromosomes, to the sex recorded during the phenotyping, (3) we compared relatedness among all individuals as determined by pedigree records to estimated relatedness using the genotype data. When the correspondence between pedigree and genetic relatedness was lower than expected for F_2_ individuals, we examined mendalization errors for all F_1_ mothers and fathers to identify the true parent, (4) we performed principal component analysis to reveal family structure, (5) for the 20 F_0_ rats that were also used for Whole genome sequencing (WGS), we compared the genotypes obtained from WGS to those obtained from the low-pass genotyping. Out of 538 samples, these steps identified 15 problematic samples. For 8 samples, the problems could be confidently resolved; however, we chose to exclude the remaining 7 samples from further analyses.

To obtain our final set of autosomal variants, a final round of variant filtering was performed by first removing SNPs with minor allele frequency (MAF) < 0.005 (SNPs with low MAF are not expected given the small number of F_0_ founders). We then removed SNPs with a Mendelian error rate < 0.075. Finally we removed SNPs with a Hardy Weinberg Error rate p < 1×10^−10^. The number of SNPs excluded by each filter are as follows: 15,879 excluded due to low MAF, 174,063 excluded due to high Mendelian Error Rate, and 55,491 excluded due to excessive HWE. After applying these filters, we were left with 4,425,349 SNPs.

### Phenotype data pre-processing and genetic analysis

Because the phenotype data were not always normally distributed, we quantile normalized the phenotypes prior to GWAS. To account for sex differences, we performed the quantile normalizations separately for males and females and then pooled the resulting values. We examined the effect of age on each phenotype using linear regression and found that age did not explain a significant proportion of variance. GWAS analysis was performed using the MLMA-LOCO algorithm of GCTA software (Yang et al., 2011) which employed a genetic relatedness matrix (GRM) and the Leave One Chromosome Out (LOCO) method to avoid proximal contamination (Cheng et al., 2013). SNP heritability estimates were also obtained with GCTA using the REML method.

Significance thresholds (alpha = 0.05) were calculated using permutation in which no GRM was used. Because the phenotype values are quantile normalized, we did not need to perform a separate permutation for each trait (Cheng & Palmer, 2013), however, given the variable sample sizes, we did perform two permutations, one for phenotypes with n>300 and one for phenotypes with n<300. For traits with sample sizes greater than 300, the significance threshold was *−log*_10_(*p*) = 6.664 and for traits with sample sizes less than 300, the threshold was slightly lower: *−log*_10_(*p*) = 6.254.

To identify QTLs, we scanned each chromosome to determine if there was at least one SNP that exceeded the permutation-derived threshold, which was supported by a second SNP within 0.5 Mb that had a p-value that was within 2 *− log*_10_(*p*) units of the index SNP. Other QTLs on the same chromosome were tested to ensure that they were conditionally independent of the first. To establish conditional independence, we used the top SNP from the first QTL as a covariate and performed a second GWAS of the chromosome in question. If the resulting GWAS had an additional SNP with a p-value that exceeded our permutation-derived threshold, it was considered to be a second, independent locus. This process was repeated (including all previously significant SNPs as covariates), until no more QTLs were detected on a given chromosome. Intervals for each identified QTL were determined by identifying all markers that had a high correlation with the peak marker >0.6 (*r*^2^ = 0.6). We performed variant annotation for all SNPs within a QTL interval using SNPEff (Cingolani et al., 2012).

## Results

### Phenotyping

Adult F_2_ animals were initially tested for their EL in a novel environment, as typically performed in the bHR and bLR selectively bred lines (e.g., F_0_s) as part of their selection criterion, followed by EPM and PavCA testing. The OF was the only test used to evaluate the adolescent F_2_ animals. Both the EPM and OF tests contain measures that reflect EL and anxiety-like behaviors. All measures were checked for possible outliers using the criteria: +/− 4 SDs from the mean; no values were removed. **Figures 1,2** and **3** contain frequency distributions of these behaviors prior to quantile normalization of the data. The lateral, rearing, and total locomotor score traits comprise EL in a novel environment (**Figure 1.A-C**). The EPM traits include distance traveled, percent time in open arms, time immobile, and number of fecal boli (**Figure.1D-F, and Figure 2.A**). OF traits include distance traveled, percent time in center, time immobile, and number of fecal boli (**Figure 1.G-I and Figure 2.B**). The PavCA traits include the PCA index from the last two test days (6 and 7) alone and the two days combined, as well as the three metrics used on each day to calculate the PCA index: response bias, probability difference, and latency score (**Figure 3.A-I**). Summary statistics for the trait data are reported in **Supplemental Table 1**.

**Figure 1.**
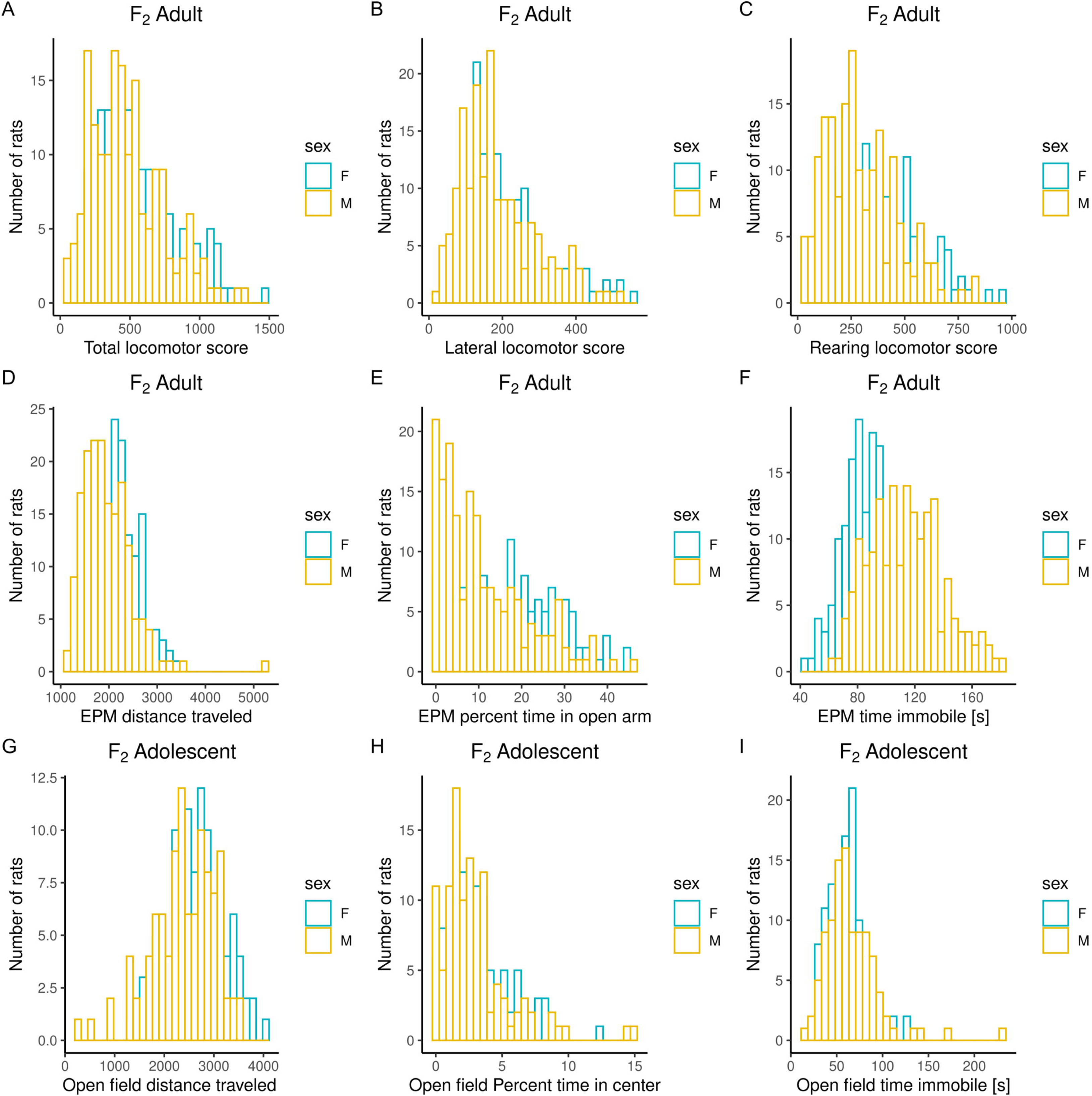
Frequency histograms showing the distribution of Locomotor Response to a Novel Environment, EPM, and OF traits in F_2_ males and females prior to quantile normalization of the data. **Figure 1.A-C**. Measures from Locomotor Response to a Novel Environment test: lateral, rearing, and total locomotor score. **Figure 1.D-F**. Measures from Elevated Plus Maze test: distance traveled, percent time in open arms, time immobile. **Figure 1.G-I**. Measures from Open Field test: distance traveled, percent time in center and time immobile. Summary statistics for the trait data are reported in **Supplemental Table 1**.

**Figure 2.**
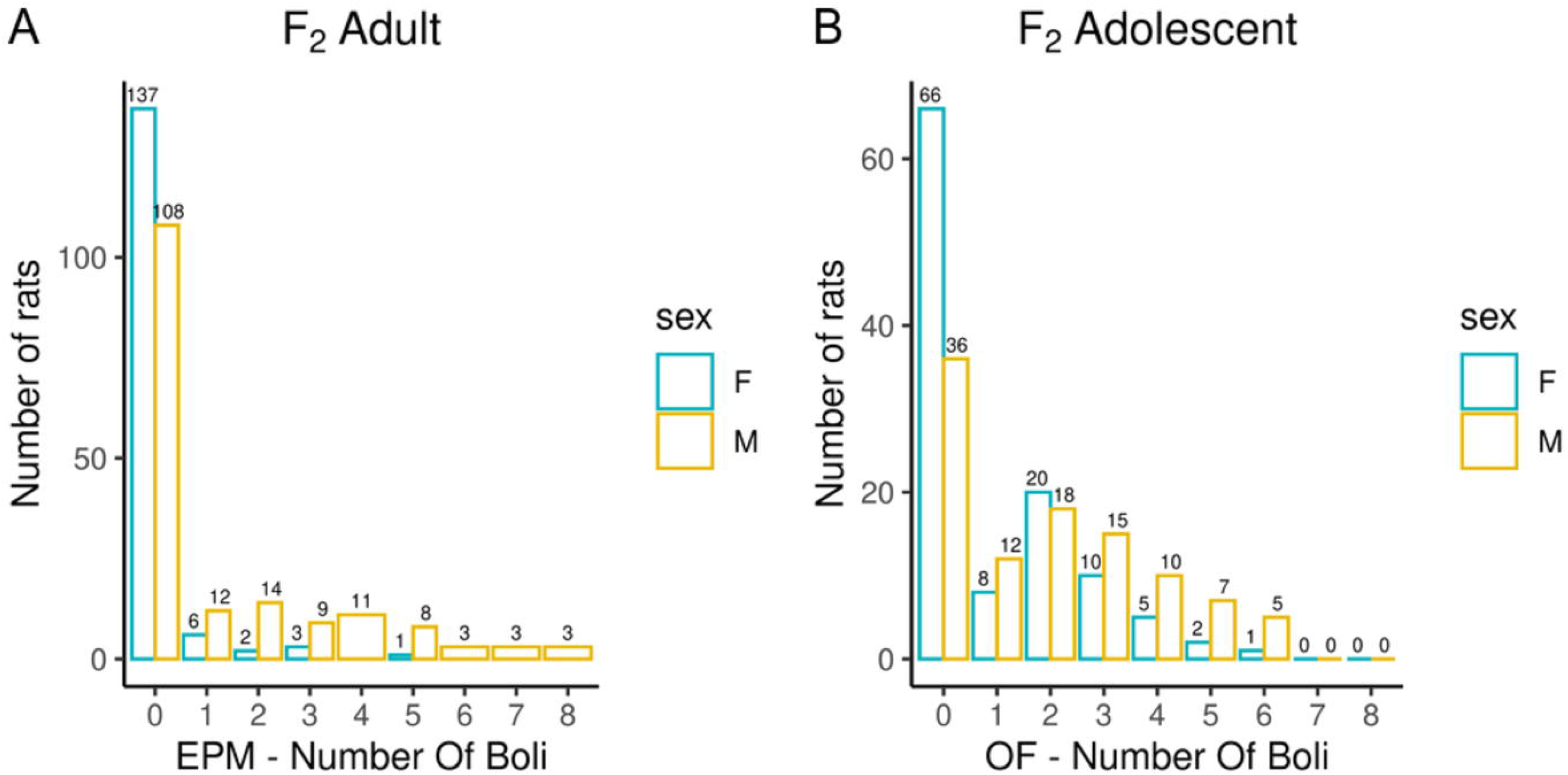
Number of fecal boli counts during EPM and OF experiments. Bar chart depicting the number of fecal boli counts colored by sex during EPM (**Figure 2.A**) and OF experiments (**Figure 2.B**). Summary statistics for the trait data are reported in **Supplemental Table 1**.

**Figure 3.**
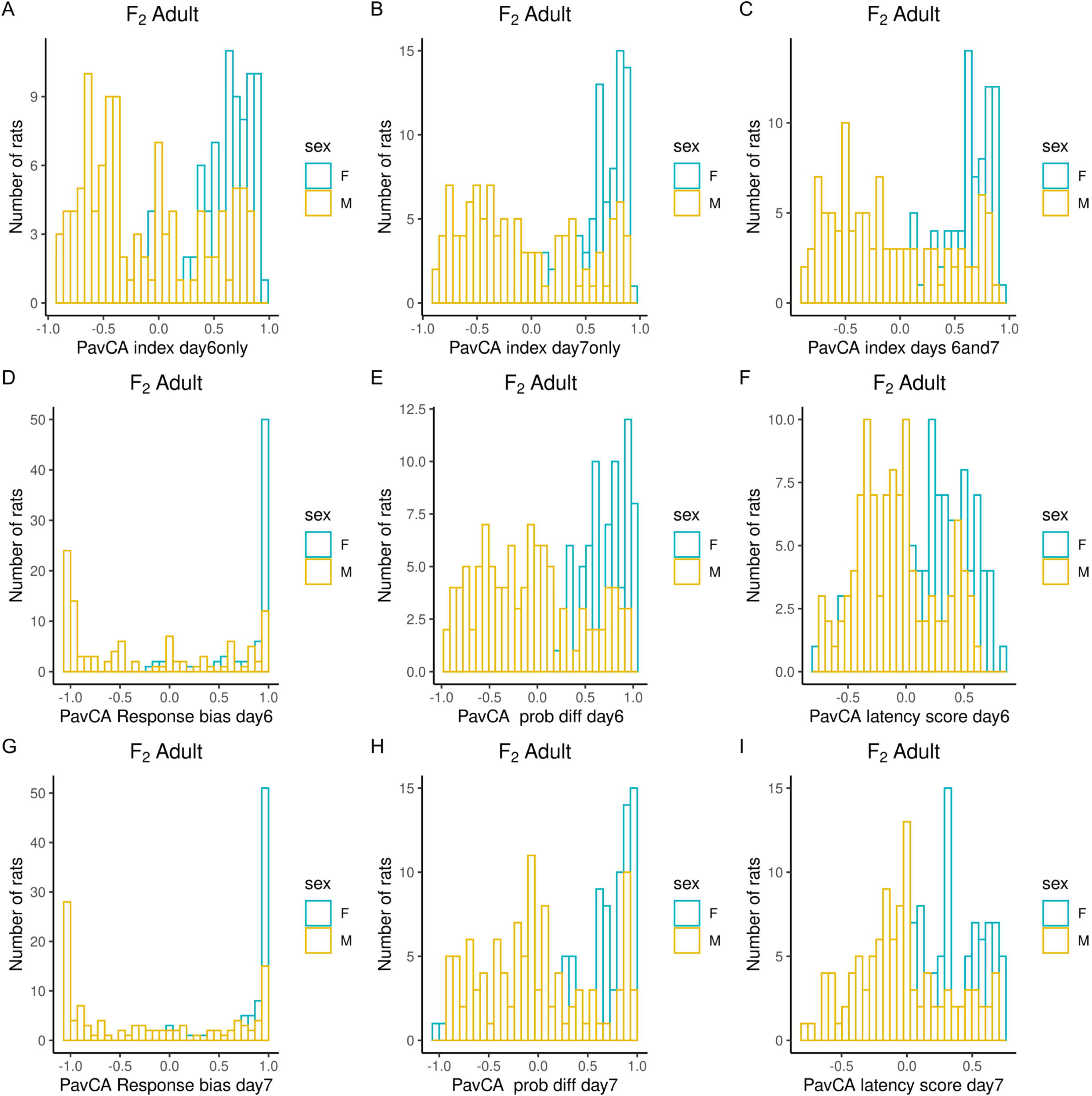
Frequency histograms showing the distribution of PavCA traits in F_2_ males and females prior to quantile normalization of the data. **Figure 3.A-I**. PCA index from the last two test days (6 and 7) alone and the two days combined, as well as the three metrics used on each day to calculate the PCA index: response bias, probability difference, and latency score. Summary statistics for the trait data are reported in **Supplemental Table 1**.

### Genotyping

Using the procedure described in the Methods section, we obtained genotypes for 4,425,349 SNPs on the autosomes. The density of the SNPs across each chromosome are depicted in **Figure 4.A**, SNP MAF distributions are in **Figure 4.B**, and the Linkage Disequilibrium (LD) decay on chromosome 20 is depicted in **Figure 4.C**.

**Figure 4.**
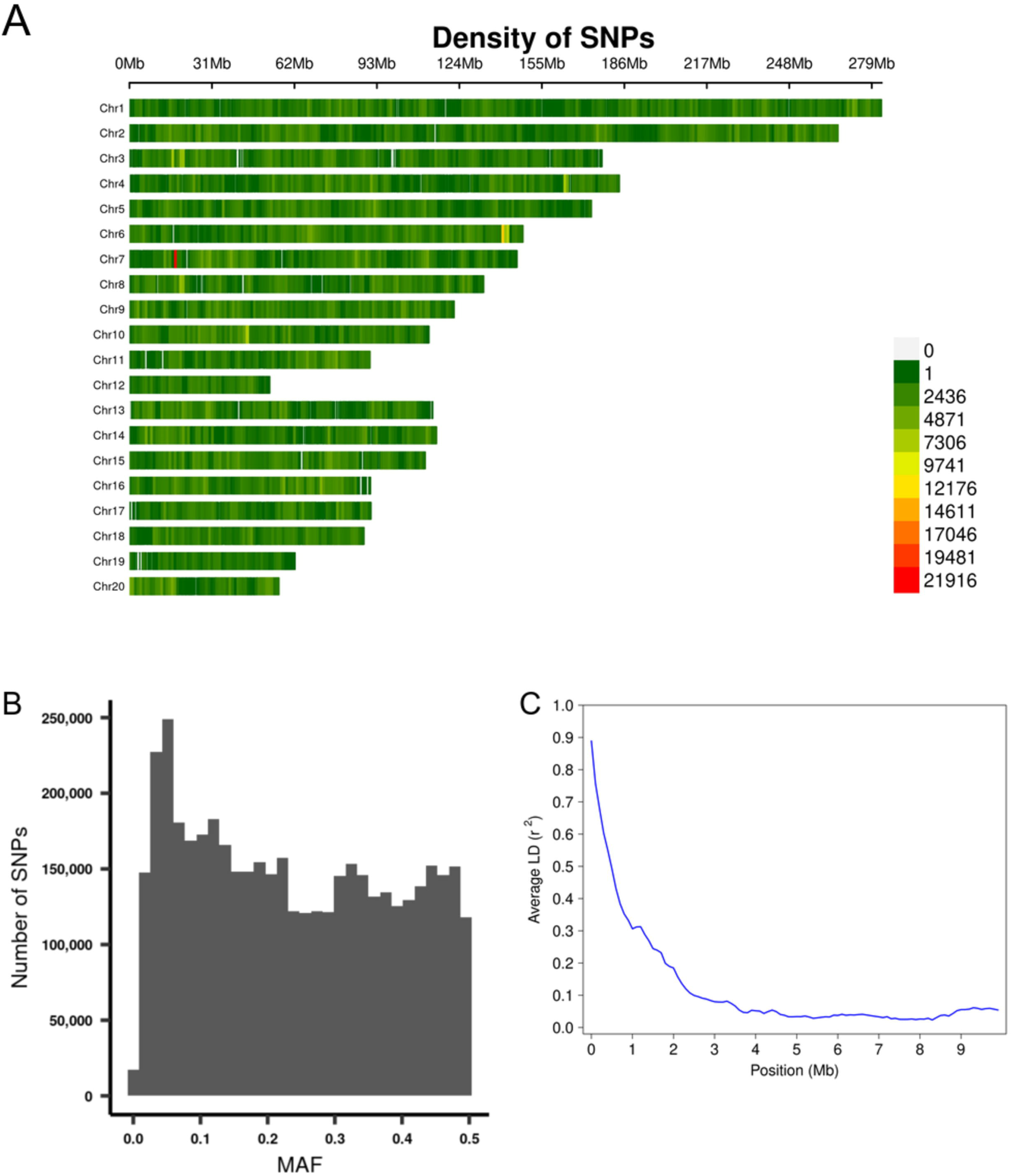
Summary of genotyping data. **Figure 4.A.** The density of the SNPs across each chromosome. **Figure 4.B.** MAF distribution of the SNPs. **Figure 4.C.** Plot of Linkage Disequilibrium (LD) decay on chromosome 20.

### SNP heritability

The SNP heritability estimates for all the phenotypes ranged from 0.06 to 0.79 (**Table 2**). Across tests, a pattern emerged relating to measures of EL, emotionality, and Pavlovian Conditioning.

**Table 2.** SNP heritability estimates. SNP *h^2^*: SNP-based heritability; SE: standard error

| Behavioral Test | Trait | SNP $h^2 \pm SE$ | P value |
| --- | --- | --- | --- |
| Locomotor Response to a Novel Environment | Total locomotor score | 0.795 ± 0.087 | 0 |
| | Lateral locomotor score | 0.796 ± 0.088 | 0 |
| | Rearing locomotor score | 0.77 ± 0.09 | 0.00E+00 |
| Elevated Plus Maze | EPM boli | 0.312 ± 0.093 | 4.25E-11 |
| | EPM distance traveled | 0.422 ± 0.113 | 1.20E-07 |
| | EPM percent time in open arm | 0.22 ± 0.088 | 1.37E-05 |
| | EPM time immobile [s] | 0.178 ± 0.106 | 5.91E-02 |
| Open Field | Open field boli | 0.429 ± 0.125 | 2.71E-09 |
| | Open field distance traveled | 0.639 ± 0.138 | 6.86E-09 |
| | Open field Percent time in center | 0.137 ± 0.115 | 8.96E-02 |
| | Open field time immobile [s] | 0.426 ± 0.135 | 3.41E-08 |
| Pavlovian Conditioned Approach | PavCA index day6only | 0.132 ± 0.116 | 1.04E-01 |
| | PavCA index day7only | 0.223 ± 0.135 | 2.47E-02 |
| | PavCA index days 6and7 | 0.26 ± 0.141 | 1.69E-02 |
| | PavCA prob diff day6 | 0.06 ± 0.098 | 2.68E-01 |
| | PavCA prob diff day7 | 0.181 ± 0.127 | 4.61E-02 |
| | PavCA response bias day6 | 0.237 ± 0.125 | 4.98E-03 |
| | PavCA response bias day7 | 0.318 ± 0.143 | 1.42E-03 |
| | PavCA latency score day6 | 0.154 ± 0.123 | 8.58E-02 |
| | PavCA latency score day7 | 0.253 ± 0.139 | 1.57E-02 |

#### EL

The EL measures revealed notably high heritability estimates across multiple tests. The highest heritability measures (>0.7) were seen across all 3 facets of EL in the primary screening test. Although the standard error estimates were higher, the “distance traveled” phenotypes for both the EPM and OF also exhibited high SNP heritability (0.4 to 0.6).

#### Emotionality

Two indices of emotionality exhibited high heritability. The first was OF Immobility (>0.4), a measure of anxiety-like behavior. The other is fecal boli counts measured during both EPM and OF. This represents another index of emotionality or anxiety-like behavior and was notably high in the OF (>0.4). EPM percent time in open arms, a classic measure of anxiety-like behavior, showed significant but lower SNP heritability (0.22).

#### Pavlovian Conditioning

The lowest SNP heritability estimates were found with the PavCA variables, although the response bias scores exhibited significant heritability (0.24-0.32).

### Phenotypic and genetic correlations

We observed a number of strong phenotypic and genetic correlations among traits measured in the adult F_2_ animals (**Figure 5**). In particular, there are strong correlations for the multiple measures derived from each test: locomotor response to a novel environment, EPM, and PavCA. There are also high phenotypic and genetic correlations for measures related to EL across tests, such as distance traveled in the EPM with the lateral, rearing, and total locomotor score traits. Although PavCA traits exhibit low SNP heritability, they are strongly correlated with EL traits, for which the genetic correlations are especially strong. **Table 3** shows how each of the PavCA measures, the combined index trait as well as the three traits that comprise the index (i.e., response bias, probability difference, and latency score) are highly correlated with the total locomotor score and the two traits that comprise this EL trait (i.e., lateral and rearing locomotor score).

**Figure 5.**
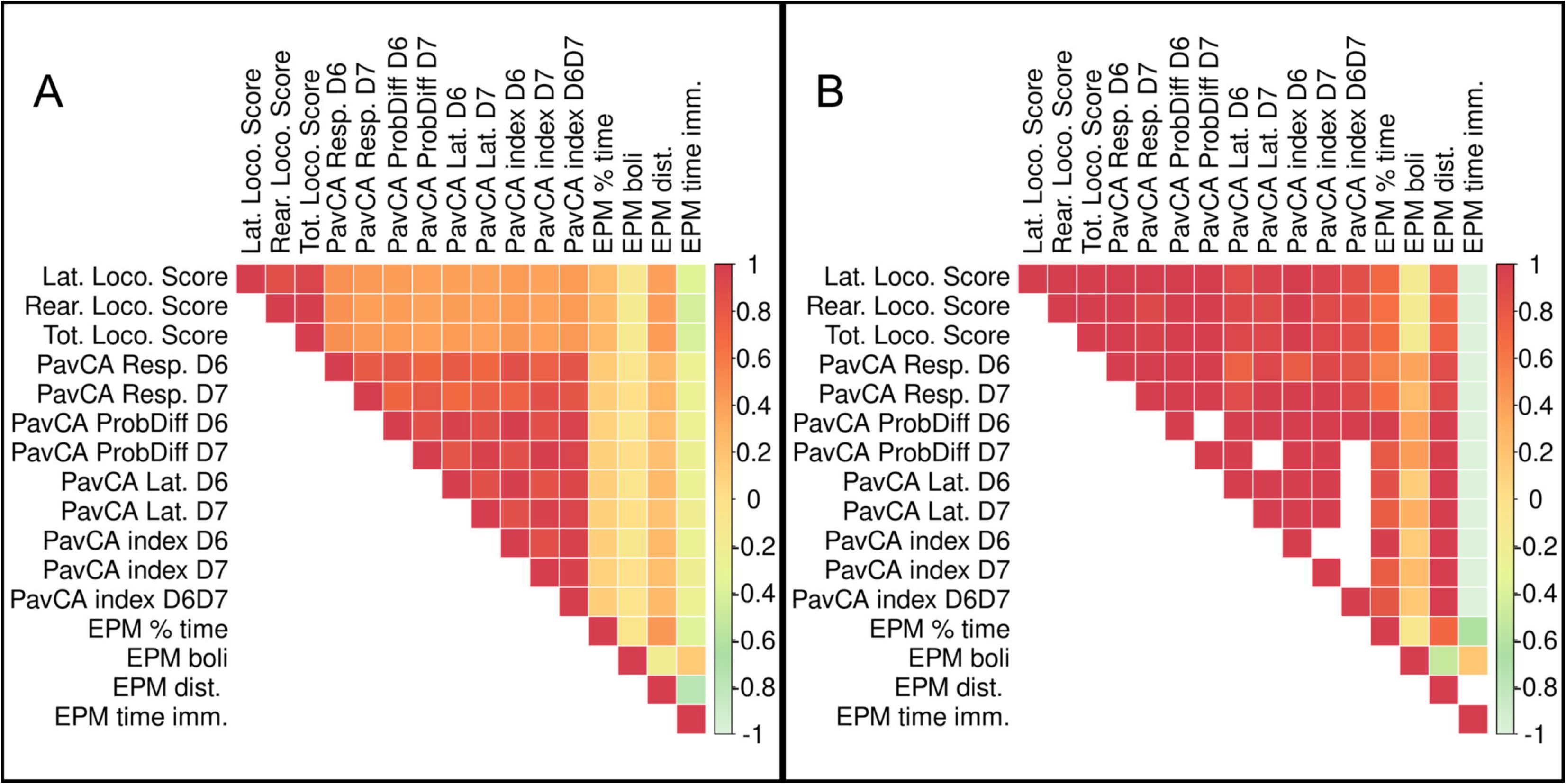
Phenotypic and genetic correlations between EPM, PavCA, and Locomotor score traits. Phenotypic and genetic correlations between EPM, PavCA, and Locomotor score (i.e, LocoScore). Phenotypic correlations are depicted in **panel A**, genetic correlations are in **panel B**. We were unable to calculate genetic correlation estimates for certain measures because the REML estimation of genetic correlation using gcta failed to converge. These are represented by blank squares. Phenotypic and genetic correlation estimates as well as the p-values are listed in **Supplemental table 2**.

**Table 3.**
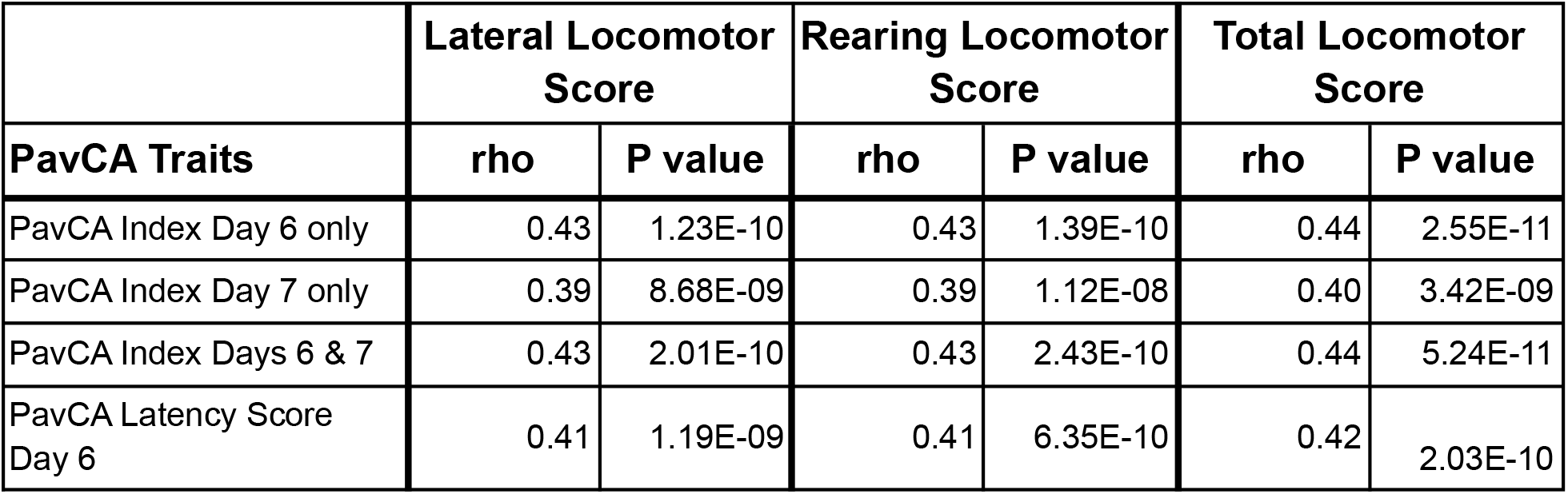
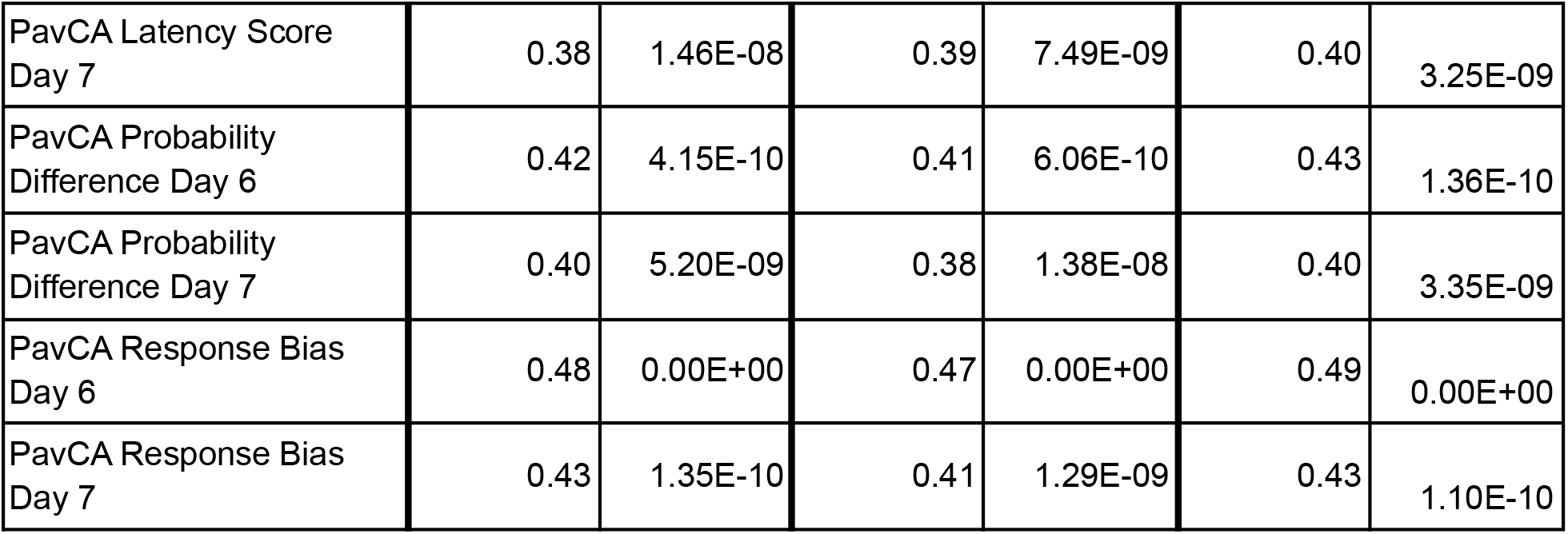
Phenotypic correlations between Locomotor Score and PavCA traits. rho: Spearman correlation.

The adolescent F_2_ animals were only tested in the OF and the correlations among the OF-related traits are depicted in **Figure 6**. The strongest positive correlations, with the genetic stronger than the phenotypic, are between the distance traveled and the percent time in the center of the OF. The strongest negative correlations, phenotypic and genetic, are between the distance traveled and the time immobile.

**Figure 6.**
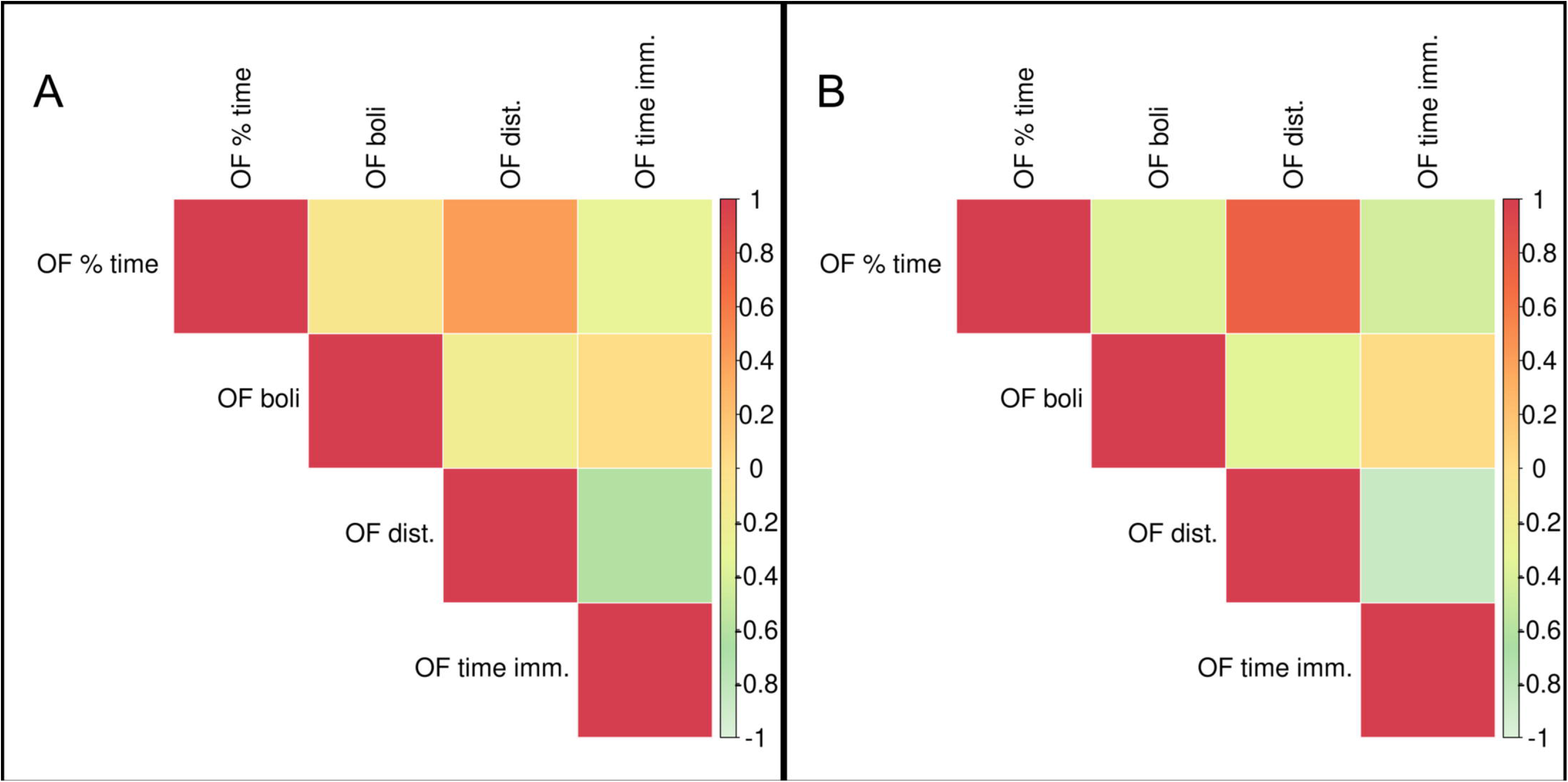
Phenotypic and genetic correlations between OF traits. Phenotypic correlations are depicted in **panel A**, genetic correlations are in **panel B**. Phenotypic and genetic correlation estimates as well as the p-values are listed in **Supplemental table 2**.

Since genetic correlations do not require both traits to be measured in the same individuals, they were calculated for the phenotypic traits in the adolescent versus adult F_2_ animals (**Figure 7**). EL traits among the three tests OF, locomotor response to a novel environment, and EPM exhibit the strongest genetic correlations, with OF time immobile highly negatively correlated with locomotor score and EPM distance traveled. PavCA traits are only significantly correlated with the OF EL distance traveled trait. Measures of anxiety or emotionality in the OF and the EPM (i.e., OF percent time in center/EPM percent time in open arms, fecal boli) are not significantly genetically correlated. However, a significant correlation exists between OF percent time in the center with two of the locomotor score traits as well as a significant negative genetic correlation between OF time immobile with EPM percent time in open arms. **Supplemental Table 2** contains genetic and phenotypic correlation estimates between all traits.

**Figure 7.**
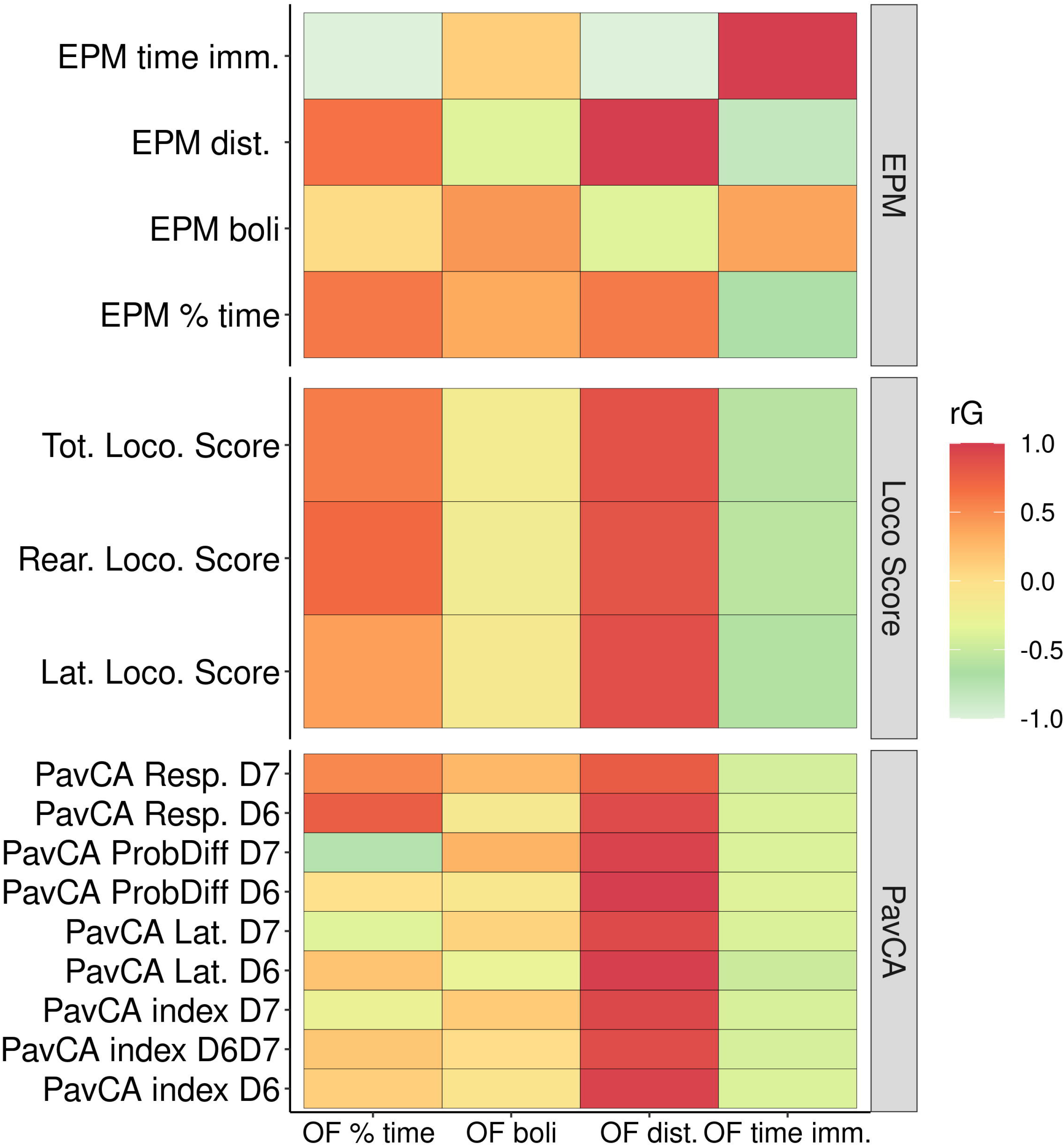
Genetic correlations between Locomotor scores, OF, EPM, and PavCA. This heat map depicts the genetic correlation estimates between Open Field measures [x-axis] against measures from EPM, Locomotor scores (i.e., LocoScore), and PavCA traits [y-axis]. Genetic correlation estimates as well as the p-values are listed in **Supplemental table 2**.

### GWAS

We discovered a total of 10 significant and conditionally independent loci for 6 behavior traits (**Table 4** and **Figure 8A**). Two of these loci appeared to be pleiotropic, meaning that the same locus was significant for multiple traits. Specifically, three traits: lateral, rearing, and total locomotor scores mapped to the same intervals on chromosome 1 at about 95 Mb and at 107 Mb. Although these loci are near one another, the peak SNPs are not in strong linkage disequilibrium (LD) and each was significant even after conditioning on the other, suggesting they represent independent loci. Another measure of activity, EPM distance traveled, also mapped to the locus on chromosome 1 at about 95 Mb. In addition, lateral locomotor score mapped to chromosome 7 at about 84 Mb. One measure of emotionality or anxiety-like behavior --OF boli-- mapped to chromosome 18 at about 23 Mb. Consistent with their lower heritability, we did not detect any significant loci for the PavCA phenotypes. Individual Manhattan plots for each trait are included in **Supplemental Figure 1**.

**Table 4.**
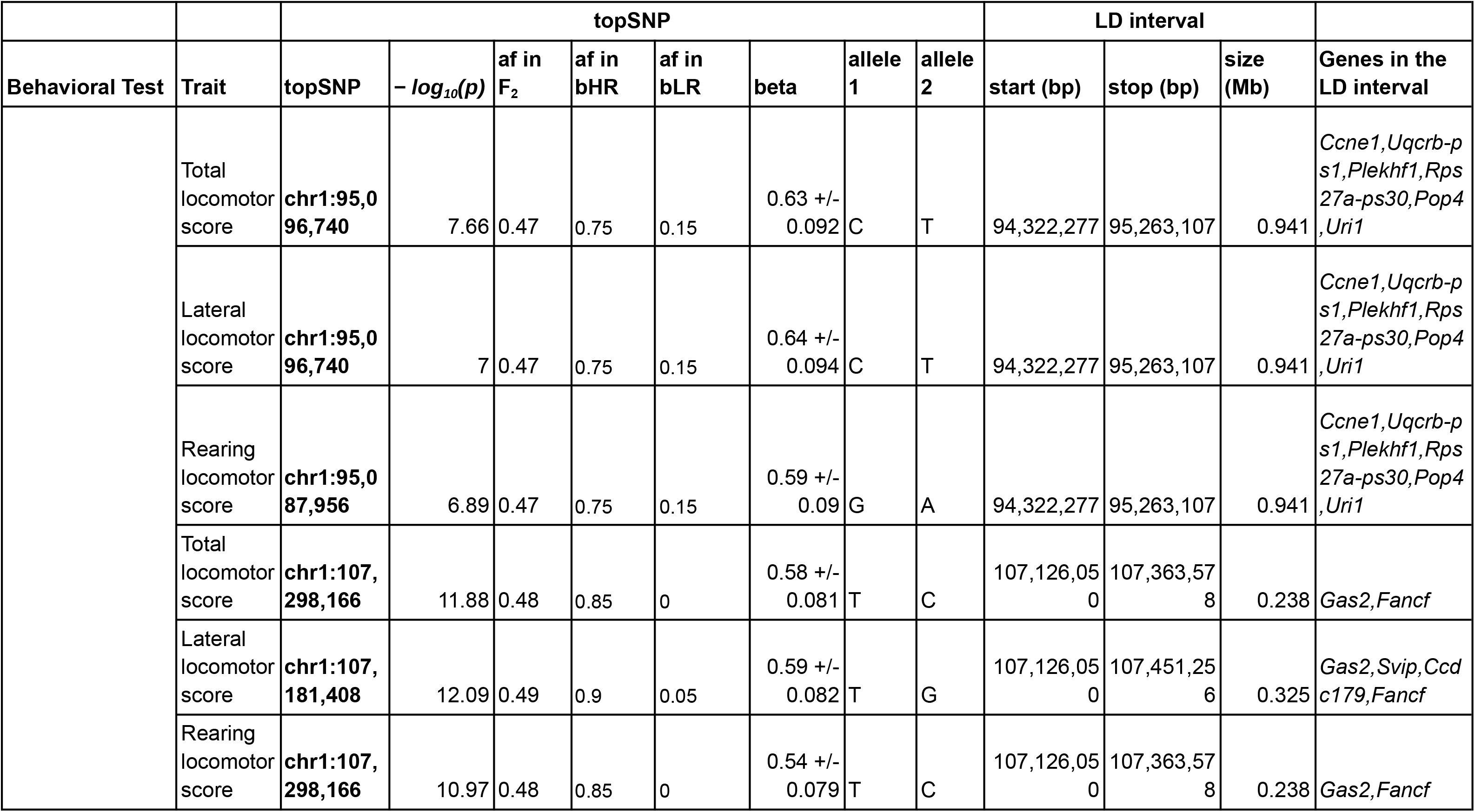
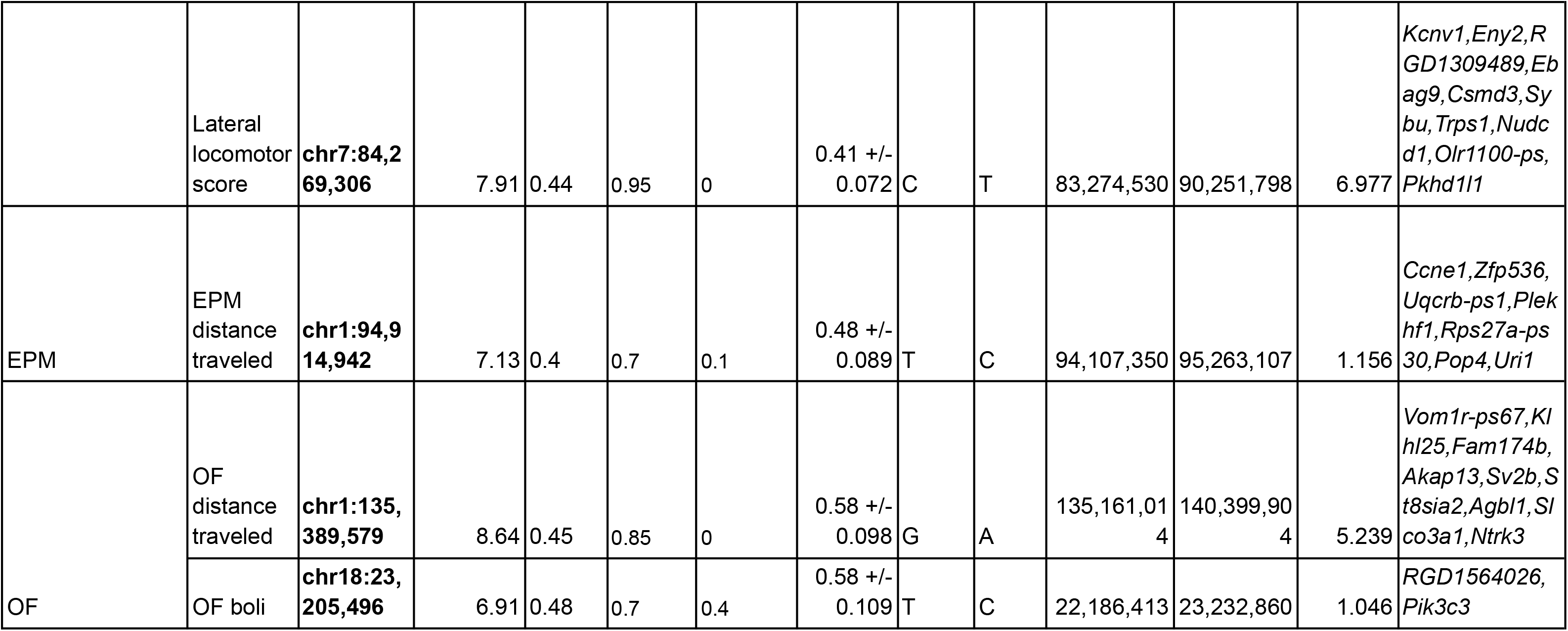
Summary of QTLs. topSNP: peak marker for the QTL; *− log*_10_(*p*): *log*_10_(*p*) for QTL peak marker; af in F_2_: Allele frequency of the effect allele in F_2_ animals; af in bHR: Allele frequency of the effect allele in 10 F_0_ bred High Responders; af in bLR: Allele frequency of the effect allele in 10 F_0_ bred Low Responders; A1: Major allele; A2: Effect allele; beta, SNP effect and standard error; start(bp) & stop (bp): physical position of the Linkage disequilibrium based QTL interval; size(Mb): size in Mb of the QTL interval.

**Figure 8.**
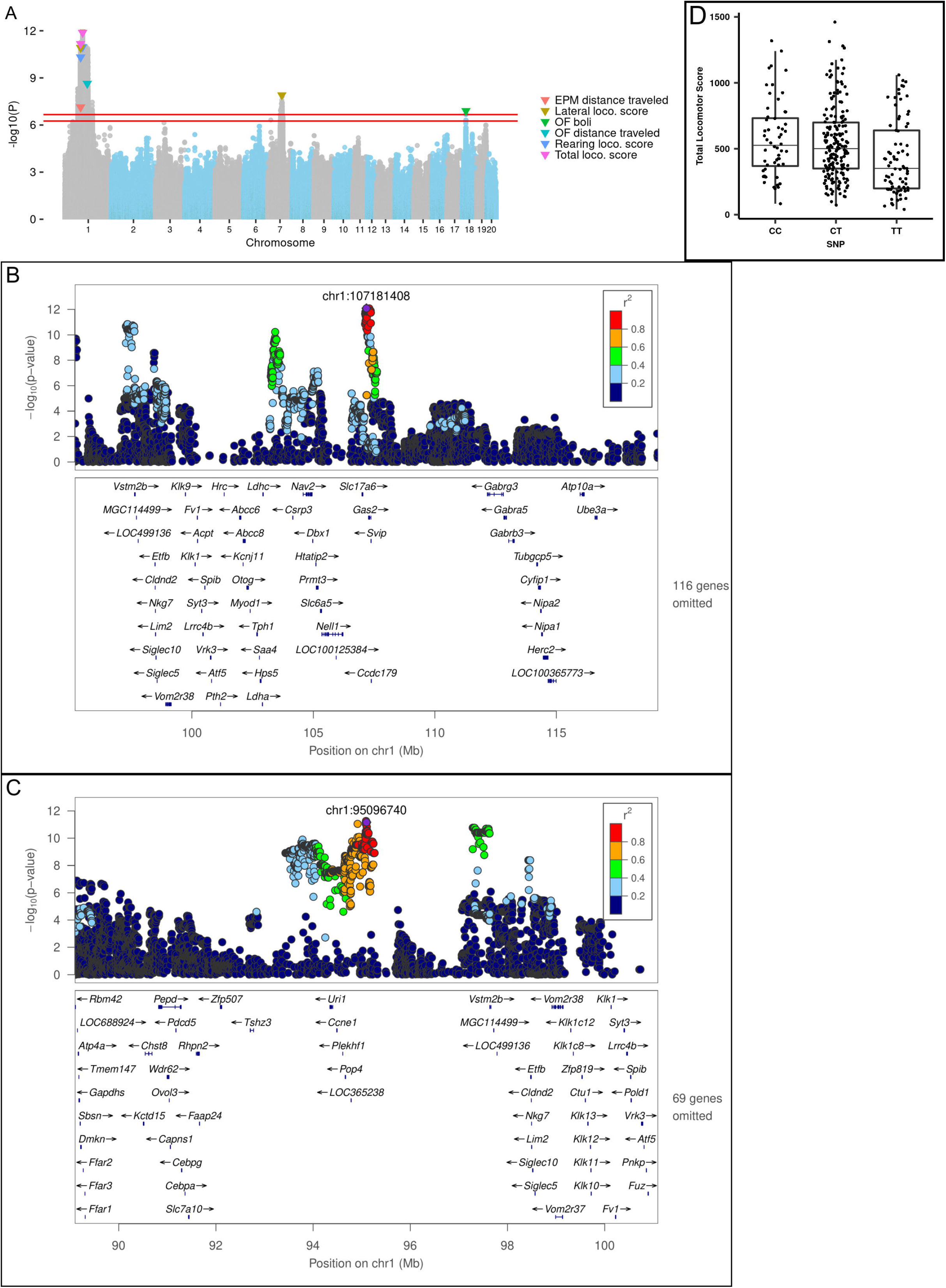
**Figure 8A. Porcupine plot for Locomotor score, EPM and OF traits**. Genome-wide association results from the GWA analysis. The chromosomal distribution of all the P-values (*− log*_10_(*p*) values) is shown, with top SNPs highlighted. The red lines show the threshold for significance. For EPM and Locomotor scores (i.e., LocoScore), the significance threshold was *− log*_10_(*p*) = 6.664 and for OF traits the threshold was *− log*_10_(*p*) = 6.254. **Figure 8B. Regional association plot for Lateral locomotor score QTL on chr1: 107Mb** **Figure 8C. Regional association plot for Total locomotor score QTL on chr1:95Mb.** X-axis shows chromosomal position. The SNPs with the lowest p-value (“top SNP”) is shown in purple, and its chromosomal position is indicated on the plot. Color of the dots indicates the level of linkage disequilibrium (LD) of each SNP with the top SNP. Genes in the region, as annotated by the RefSeq. **Figure 8D. Effect plot for Total locomotor score QTL on chr1:95Mb.** Effect plot depicting the genetic effect of the peak SNP chr1: 95 Mb with a significant association to Total Locomotor Score.

The loci identified ranged in size from 239 Kb to 6.98 Mb and contained between 2 and 10 genes. LocusZoom plots for the two loci on chromosome 1 are shown in **Figures 8B and 8C**. An effect plot depicting the genetic effect of the peak SNP at chromosome 1:95 Mb with a significant association to total locomotor score is shown in **Figure 8D**. Additional LocusZoom plots for the other loci are included in **Supplemental Figure 2**. We identified 5 potential coding variants affecting the *Plekhf1* and *Pkhd1l1* genes on chromosomes 1 and 7, respectively, that were predicted to have moderate impact and are shown in **Table 5**. **Supplemental Tables 3 and 4** contain the genotypes of the F_0_ rats at the moderate impact variants and QTL peak SNPs.

**Table 5.** Summary of putatively causal coding variants. QTL: peak marker of the QTL; SNP in LD with peak marker: Moderate impact variant in LD with peak marker of the QTL; LD r^2^ with trait topSNP: Linkage Disequilibrium correlation coefficient of moderate impact variant with peak marker; SNP change in HGVS notation; Amino acid change in HGVS notation.

| Trait | QTL | Candidate gene | SNP in LD with peak marker | LD r<sup>2</sup> with trait topSNP | SNP change | Amino acid change |
| --- | --- | --- | --- | --- | --- | --- |
| EPM distance traveled | chr1:94 Mb | *Plekhf1* | chr1:94,610,123 | 0.69 | c.706A>G | p.Ser236Gly |
| Lateral locomotor score | chr1:95 Mb | *Plekhf1* | chr1:94,610,123 | 0.65 | c.706A>G | p.Ser236Gly |
| Lateral locomotor score | chr7:84 Mb | *Pkhd1l1* | chr7:83,403,144 | 0.77 | c.716A>C | p.Lys239Thr |
| Rearing locomotor score | chr1:95 Mb | *Plekhf1* | chr1:94,610,123 | 0.65 | c.706A>G | p.Ser236Gly |
| Total locomotor score | chr1:95 Mb | *Plekhf1* | chr1:94,610,123 | 0.65 | c.706A>G | p.Ser236Gly |

## Discussion

In the current study, we probed genetic differences produced by a long-term selective breeding program by creating an F_2_ cross between two phenotypically divergent outbred rat lines and then used this F_2_ cross to map loci for the selection trait, sensation-seeking behavior, as well as several putatively correlated traits. We identified high heritability not only of EL measures but also of other traits associated with anxiety-like behavior. We also uncovered significant correlations among traits that characterize these bred lines, both at the behavioral and genetic level.

Despite the modest (for GWAS) sample size, we identified significant genetic associations for all facets of EL, OF, and EPM behaviors, but not for PavCA behavior. We also discovered that different measures of EL were influenced by pleiotropic loci, which was expected given the strong phenotypic and genetic correlations among those traits. The loci were relatively small and contained several interesting genes, including some that harbor coding variants with modest impact.

### Strong phenotypic and genetic correlations among traits

Substantial heritability exists among several traits besides EL in the F_2_ animals. We have previously identified high heritability in the total locomotor score trait of our bHR/bLR animal model, for which EL was defined only by the total measure of locomotion in a novel environment (Zhou et al., 2019). We have now expanded the definition of EL to encompass the components of the total locomotor score, both lateral and rearing locomotor score, and the distance traveled in the EPM and the OF. Our current results reveal high heritability among all these measures of EL encompassing three behavioral tests assessing EL in different environmental contexts. Significant heritability in EL is consistent with a recent report demonstrating such heritability in the genetically diverse Collaborative Cross inbred mouse model consisting of eight founder strains and 10 Collaborative Cross strains (Bailey et al., 2021). Both phenotypic and genetic correlations are also quite strong among these traits in the F_2_ animals, with anxiety-related traits showing moderate correlations, indicating a similar genetic architecture for these behaviors. Although PavCA traits exhibit low SNP heritability as has been reported for Sprague-Dawley rats (Gileta et al., 2022), they are strongly correlated with EL traits in our F_2_ animals, and the genetic correlations are especially strong. This is notable as outbred rats have not shown such a relationship between EL and sign-tracking/goal-tracking behavior (Flagel et al., 2010; Hughson et al., 2019; Robinson & Flagel, 2009). As previously suggested (Flagel et al, 2010), it appears the selective breeding co-selected the sign-tracking/goal-tracking trait along with the sensation-seeking trait, and the present findings point to the genetic features of this association.

### Genetic loci identified for EL traits and comparison with a previous study

In the current study, we used approximately 4,425,349 variants in 538 F_2_ rats for genetic mapping to identify significant QTLs for Locomotor score, EPM, and PavCA traits in the adult animals and OF traits in the adolescent animals. In a previous study (Zhou et al., 2019) that used some of the same F_2_ rats but a much smaller set of just 416 markers that only included exonic SNPs, we identified loci on chromosomes 1, 2, 3, 7 and 18 for total locomotor score (EL in Zhou et al., 2019). While there is broad concordance between the two studies, only the top SNP from chromosome 7 for the QTL from Zhou et al., 2019 lies within the LD interval for the QTL we detected on chromosome 7 in the current study for lateral locomotor score. **Supplemental Table 5** shows the relationship between Zhou et al., 2019 and results from the current study. There are multiple reasons that could explain this lack of overlap, including that the previous study employed fewer markers, which can impact the location of associations. We also performed somewhat different analyses (our study used MLMA-LOCO in GCTA and included a random term to account for familial relationships whereas QTL analyses in the previous study was performed using R/qtl), which could have led to the divergent results between the two studies.

Below, we discuss several of the loci identified in this analysis. Parallel analyses are underway to determine whether the observed variants are associated with differences in gene expression in the brain of the bHR, bLR, or the F_2_ cross, and how expression levels might relate to behavioral traits (Hebda-Bauer, Hagenauer et al, in preparation).

### Identification of a missense variant in *Plekhf1* gene on chromosome 1:95 Mb QTL for EL traits

Within the locus on chromosome 1 at about 95 Mb to which the main EL traits mapped (**Figure 8C**), we identified a missense variant in the gene *Plekhf1*, which has been implicated in apoptosis and autophagy (Lin et al., 2012). Interestingly, *Plekhf1* has previously been associated with glucocorticoid receptor signaling in mice and rats. This missense variant is notable since glucocorticoid signaling, as part of the hypothalamic-pituitary-adrenal axis, is dysregulated in major depressive disorder (McEwen & Akil, 2020). Sato et al., 2008 identified *Plekhf1* as one of the genes responding to glucocorticoids in the hypothalamus of Sprague-Dawley rats. Luo et al., 2018 found that *Plekhf1* was upregulated in the hippocampus of mice with low anxiety- and low depressive-like phenotypes. Mice with increased expression of *Plekhf1* exhibited higher EL and less anxiety-like behavior (i.e, increased distance traveled in the center) in the OF test, consistent with the identification of this locus in association with EL in the current study. In a study that investigated the long-term impact of early life adversity on the ability to cope in adulthood, it was found that chronic early life stress in mice generally blunted the translational response to an adult acute stress challenge. In a previous study, *Plekhf1* was one of only 18 genes identified as being upregulated in the hippocampus by acute-swim stress in mice that had experienced early life stress (Marrocco et al., 2019).

*Plekhf1* is also differentially regulated in the mouse brain in response to various drugs of abuse (Ficek et al., 2016; Piechota et al., 2010; Wu et al., 2018). *Plekhf1* was one of 16 genes that was significantly differentially expressed in the striatum between ketamine- and saline-treated mice (Ficek et al., 2016). Ketamine, an NMDA receptor antagonist, is known for its rapid antidepressant action and its gene expression profile was found by Ficek et al (2016) to be more similar to opioids, ethanol, and nicotine than to psychostimulants. Piechota and colleagues (2010) revealed that 42 drug-responsive genes in the striatum cluster into two gene expression patterns with *Plekhf1* identified in the first sub-cluster of the second pattern that shows an early response to ethanol, morphine, and heroin. In another study, *Plekhf1* was upregulated by both morphine and heroin and is associated with psychological dependence to these drugs (Wu et al, 2018). Thus, ample evidence exists for an important role of the *Plekhf1* gene in stress-related behavioral effects, mood disorders, and substance use.

### Identification of *Fancf* and *Gas2* as potential candidate genes that may drive the chromosome 1:107 Mb QTL for EL traits

For the locus on chromosome 1 at about 107 Mb that is also associated with the EL traits, we identified two attractive candidate genes: *Fancf* and *Gas2*. *Fancf* is one of the top genes associated with a SNP differentiating motor subtypes in Parkinson’s disease in a human GWAS meta-analysis (Alfradique-Dunham et al., 2021). The *Gas2* protein family is important for physiological processes such as cytoskeletal regulation, cell cycle, apoptosis, senescence, and differentiation (Zhang et al., 2021) and pathological processes such as human brain glioma development (Zhao et al., 2021). Recently, a human GWAS revealed that *Gas2* is located in genetic loci influencing general cognitive ability, suggesting a novel connection between this gene and cognition (G. Davies et al., 2018). Thus, further evaluation of the role of *Gas2* could potentially lead to new mechanisms for sensation-seeking behavior.

### Distance traveled measures from OF and EPM map to different intervals

Through our analyses, we also found that the genetic loci for the distance traveled measures from OF and EPM do not overlap. OF distance traveled maps to chromosome 1:135 Mb, while EPM distance traveled maps to chromosome 1:95 Mb. The distance traveled in the OF and the EPM likely represent different aspects of novelty-induced locomotor activity (i.e., EL) that may also be influenced by developmental age (e.g., adolescence versus adulthood). The chromosome 1:135 Mb locus contains many genes. Several of these genes have been previously associated with neurodevelopmental disorders in humans. *Fam174b* is implicated in autism spectrum disorder (Kamien et al., 2014) and variants of the *St8sia2* gene increase susceptibility to bipolar disorder, schizophrenia, and autism (Shaw et al., 2014). *Ntrk3* is associated with eating disorders (Mercader et al., 2008), panic disorders (Armengol et al., 2002) and onset of childhood mood disorders (Feng et al., 2008). *Ntrk3*, neurotrophic tyrosine kinase receptor 3, is a member of the TRK family of tyrosine protein kinase genes and the protein product of this gene is trkC, which is preferentially expressed in brain (Feng et al., 2008). The identification of a genetic locus at chromosome 1:135 Mb for locomotion in a novel stressful environment for young animals, which contains *Ntrk3*, is consistent with our previous findings of gene expression differences in a neurotrophic factor (i.e., FGF2) between the bHR and bLR animals and how augmentation with FGF2 early in life rescues the anxiety-like phenotype in the bLRs that have low FGF2 expression (Turner et al., 2011).

### Identification and replication of *Pkhd1l1 and Trhr* for locomotor activity measure

We identified a missense variant within the *Pkhd1l1* gene for the chromosome 7:84 Mb locus for lateral locomotor score in response to novelty. *Pkhd1l1* has been associated with bipolar disorder in humans (Forstner et al., 2020). This locus also contains the gene *Trhr* which is an important component of the hypothalamic-pituitary-thyroid axis and regulates anxiety- and depressive-like behavior (Choi et al., 2015; Pekary et al., 2015; Zeng et al., 2007). Our group has previously identified both *Trhr* and *Pkhd1l1* to be differentially expressed in hippocampus of bHR/bLR rats and has shown that *Trhr* correlates positively with locomotor activity (Birt et al., 2021). Both *Trhr* and *Pkhd1l1* were also found to be 4.5 times more likely to be located within a QTL for EL (LOD>4) identified in our previous exome sequencing study (Zhou et al., 2019).

### *Pik3c3* within the chromosome 18:23 Mb locus for OF fecal boli measure

The locus on chromosome 18 at about 23 Mb that was associated with OF fecal boli contains the gene *Pik3c3* which is involved in phosphoinositide lipid metabolism, a potential target for the therapeutic effect of lithium in bipolar disorder. A promoter variant of this gene has been identified in humans with bipolar disorder and schizophrenia (Stopkova et al., 2004). Fecal boli are released as part of the stress response and are considered a measure of emotional reactivity or anxiety-like behavior; thus, the identification of this locus from adolescent animals suggests the potential developmental contribution of this locus to emotional reactivity and vulnerability to adverse stress-related effects. The selectively bred bLR rats exhibit significantly more fecal boli during stress- and anxiety-like tests compared to the bHR rats (unpublished data), which is consistent with their internalizing phenotype comprising anxiety- and depressive-like behaviors. Thus, identification of a significant locus for fecal boli in young F_2_ animals lends support for a genetic basis for this trait.

### Allele frequency for QTL topSNP in the F_0_ bHR and bLR rats

We looked at the allele frequency for each QTL topSNP in the 20 F_0_ bHR and bLR rats. (**Table 4**) As expected, the HR alleles were always associated with higher activity. For most bLR animals, the alternative allele frequency was zero, showing that protracted selection sometimes led to fixation (since we only sequenced 10 bHR rats, it is possible that the alternate allele still exists at low frequency in the bHR). For several other alleles, there was a large difference in allele frequencies, but both alleles were found in both populations. This observation is consistent with a previous study (Phillips et al., 2002) in which reverse selection revealed a lack of fixation of selection relevant alleles even after a protracted period of selection.

## Conclusions

In conclusion, we have demonstrated strong heritability in the sensation-seeking trait, depicted by the EL measures, through the F_0_-F_1_-F_2_ cross of the bHR/bLR rat lines and discovered several genetic loci associated with complex behavior traits. Although we found strong genetic correlation between EL and PavCA measures, no significant loci for PavCA traits were identified. Remarkably, while the breeding program selected for high and low EL in a novel environment, our analyses revealed loci of relevance to different facets of anxiety, including freezing behavior in an open field and the number of fecal boli under stress. We also discovered putative pleiotropic loci that influence EL traits across different behavioral assays, thus expanding upon our previous finding of several loci tied to EL in only our main behavioral assay used for selective breeding. The strong heritability coupled with identification of several loci associated with EL and emotionality provide compelling support for this selectively bred rat model in discovering causal variants tied to elements of internalizing and externalizing behaviors inherent to psychiatric and substance use disorders. The candidate genes located in the significant loci associated with these sensation-seeking and anxiety-like traits provide avenues for further discovery into the genetic basis of stress, affect regulation, and addiction-related traits.

## Acknowledgements

NIDA U01DA043098 (HA, JZL, AP): Office of Naval Research (ONR) 00014-19-1-2149 (HA); The Hope for Depression Research Foundation (HDRF) (HA); the Pritzker Neuropsychiatric Research Consortium.

