## Supplemental figures and tables for "Genome-Wide Association Study in a Rat Model of Temperament Identifies Multiple Loci for Exploratory Locomotion and Anxiety-Like Traits"

**Supplemental Figure 1.** Manhattan plots for all traits

EPM boli

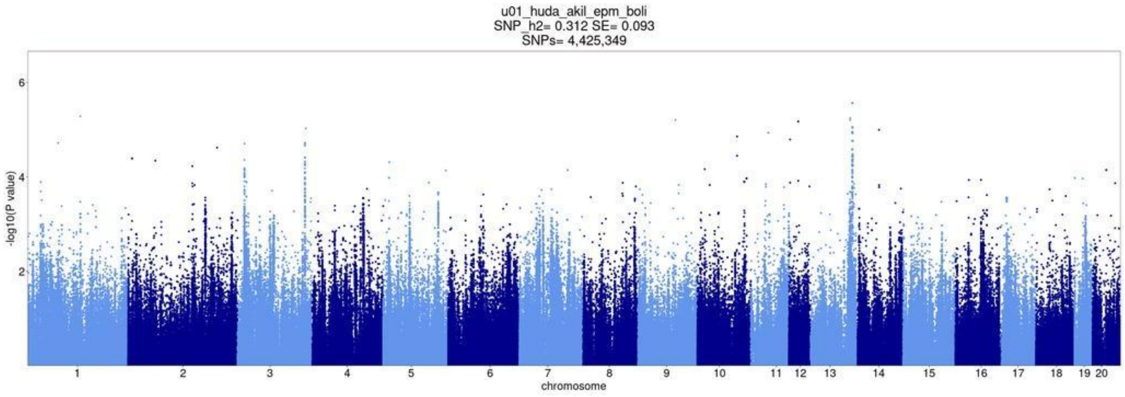


EPM distance traveled

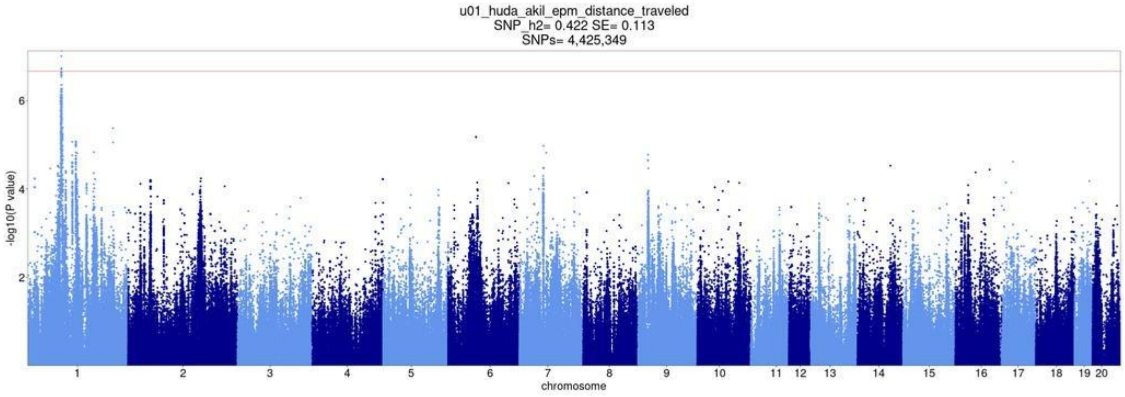


EPM percent time in open arm

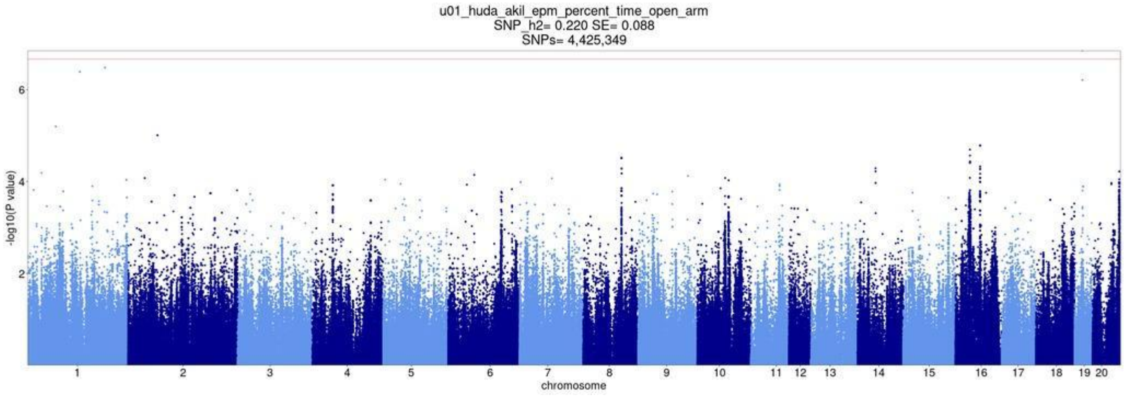

EPM time immobile [s]

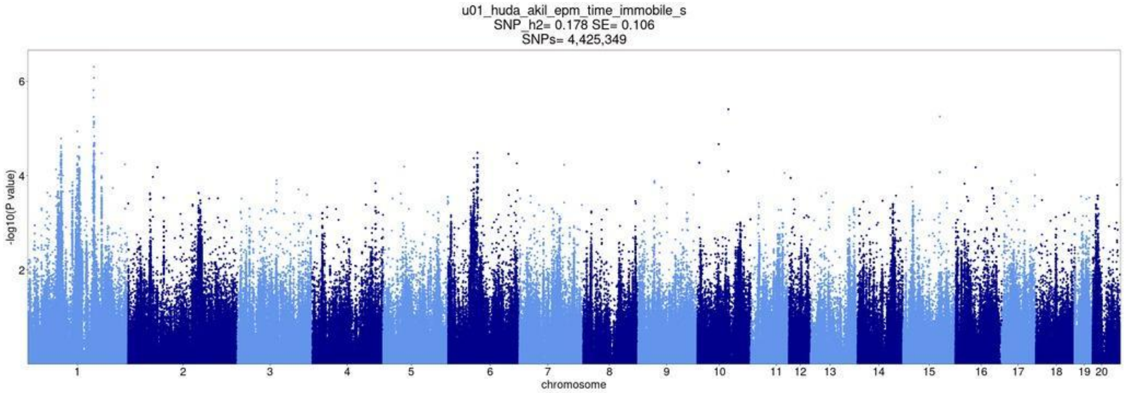


PavCA latency score day6

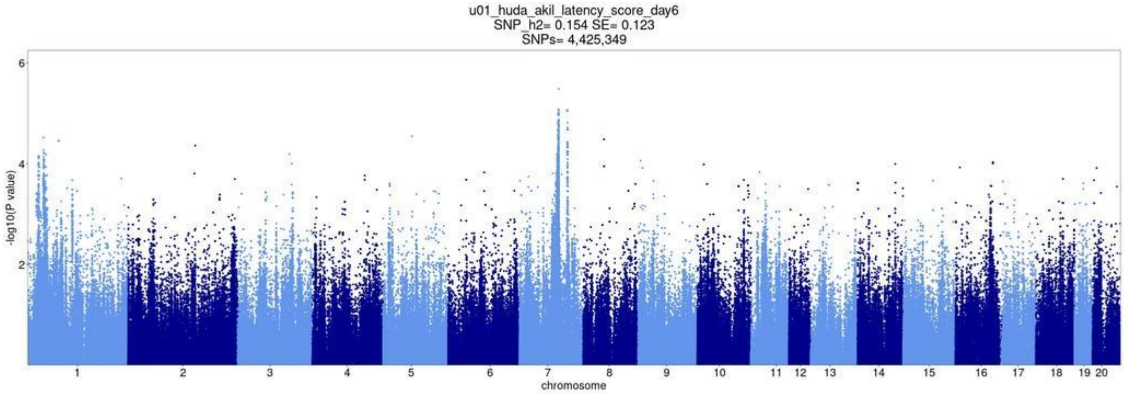


PavCA latency score day7

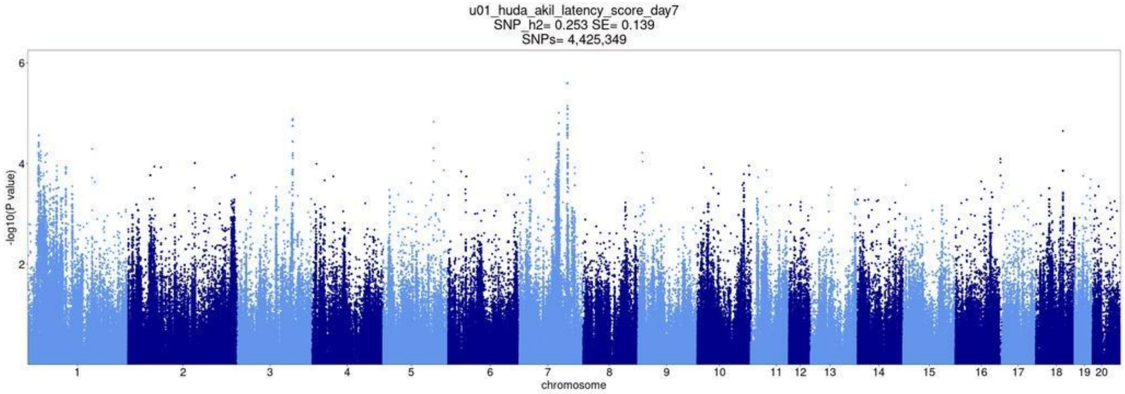

Lateral locomotor score

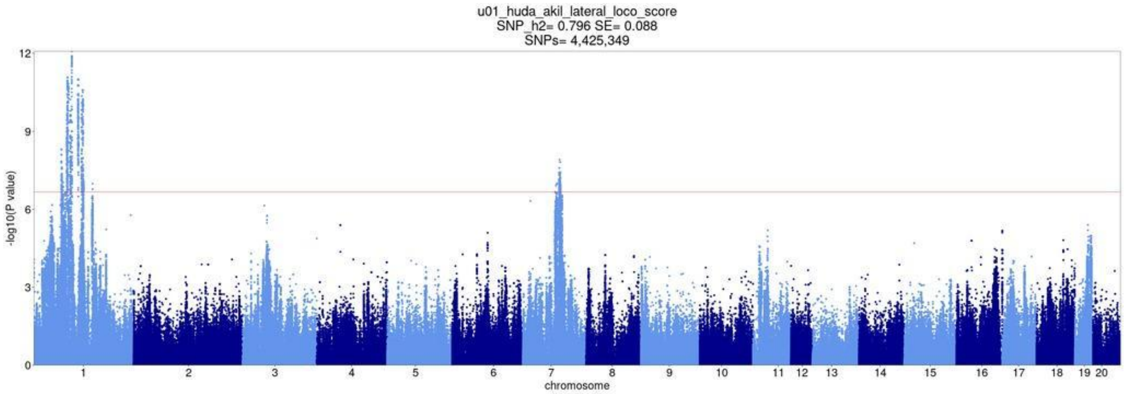


Open field boli

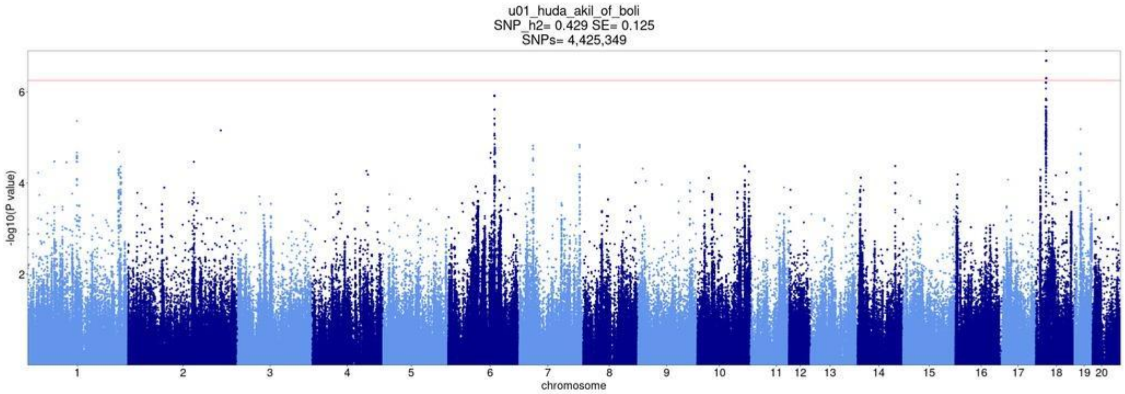


Open field distance traveled

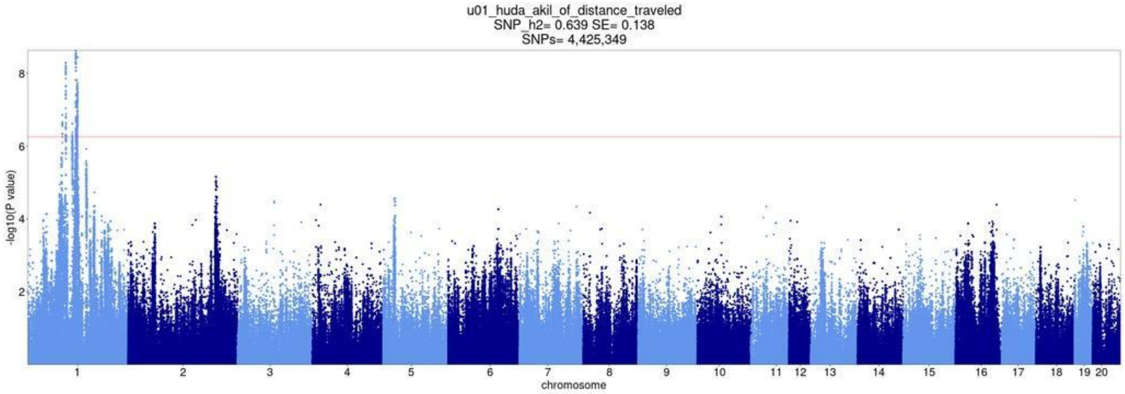

Open field Percent time in center

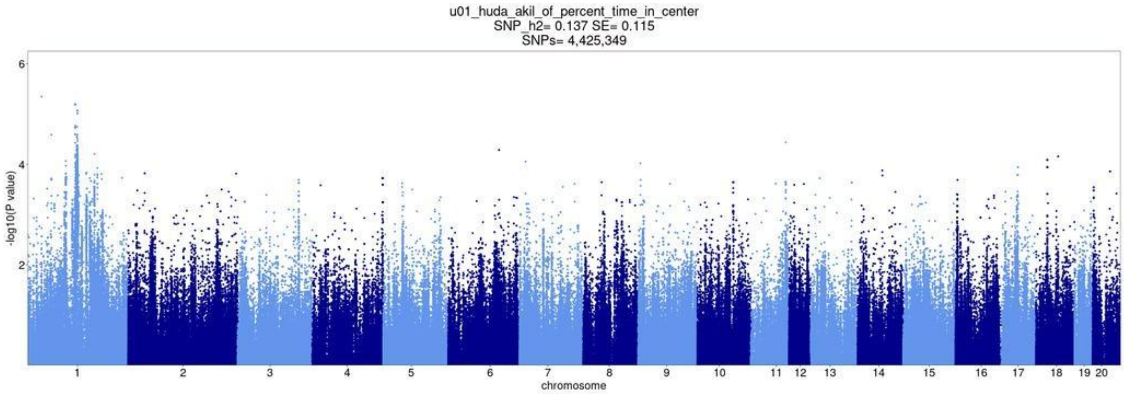


Open field time immobile [s]

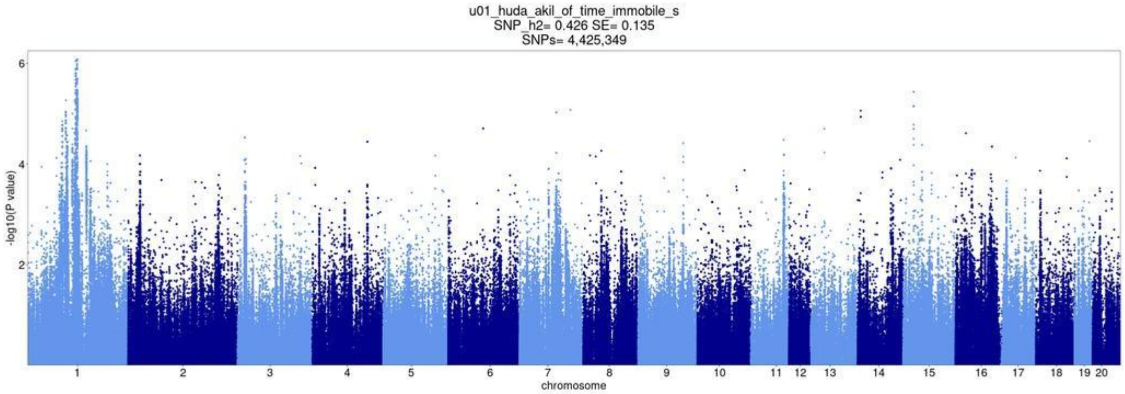


PavCA index day6only

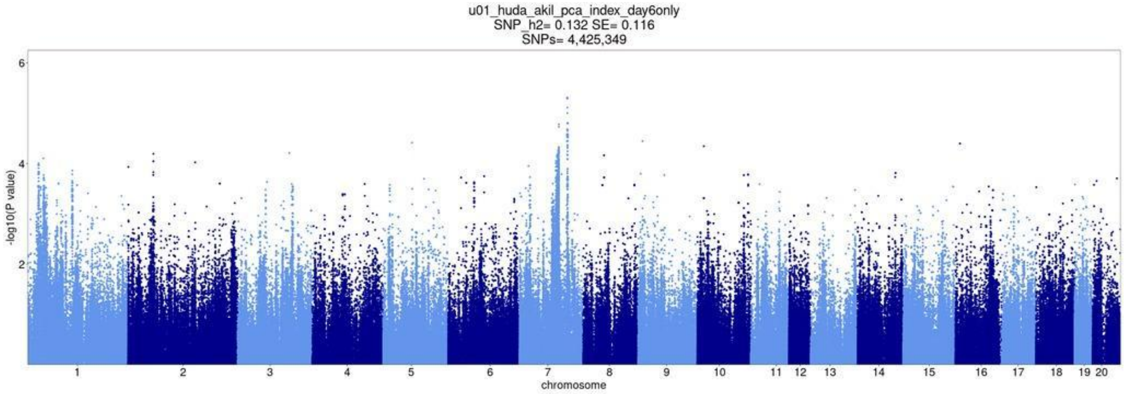

PavCA index day7only

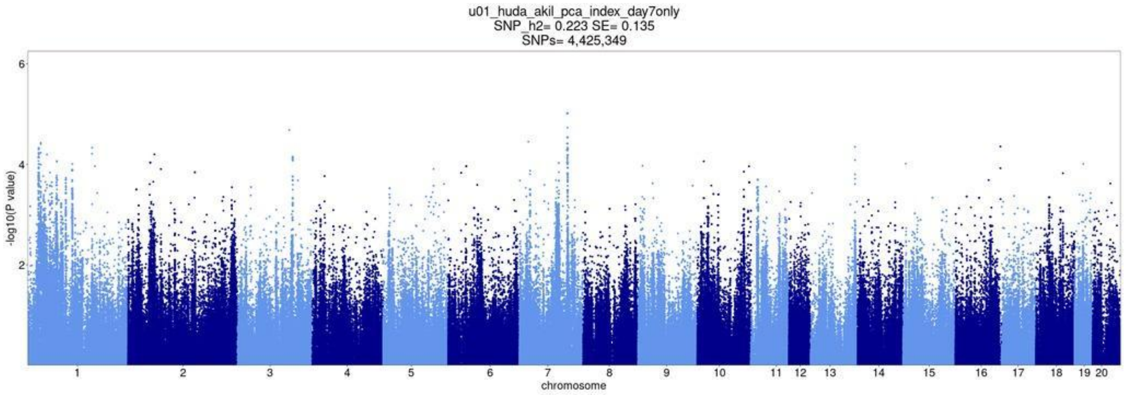


PavCA index days 6and7

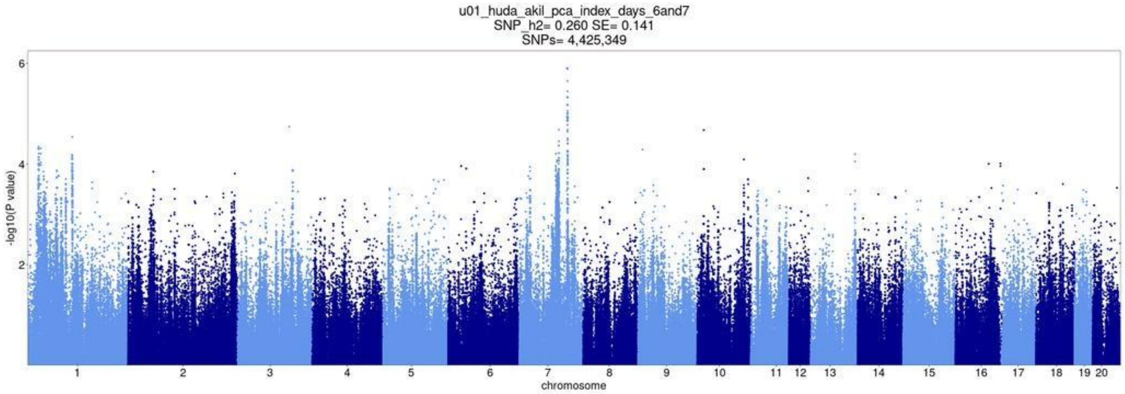


PavCA prob diff day6

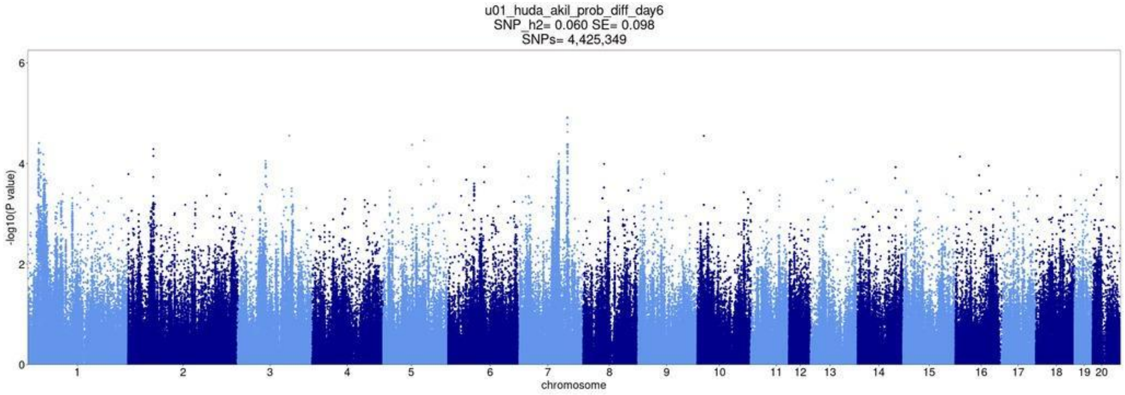

PavCA prob diff day7

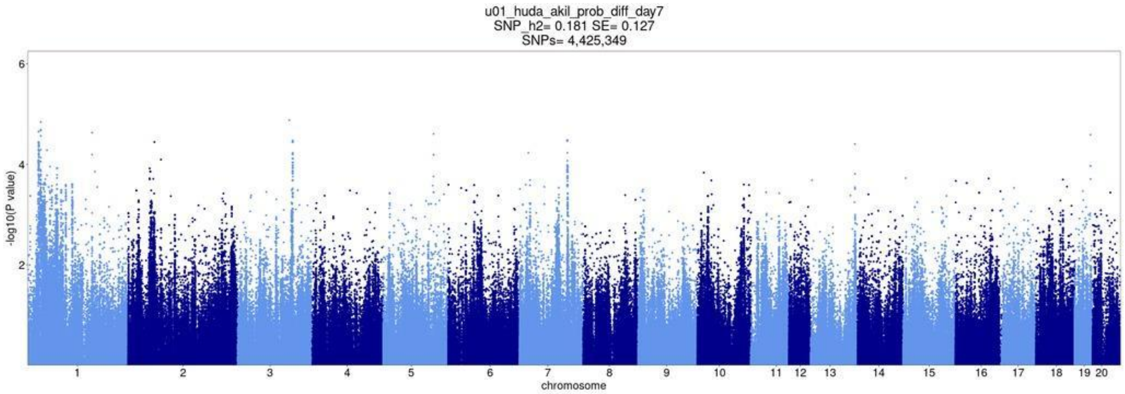


Rearing locomotor score

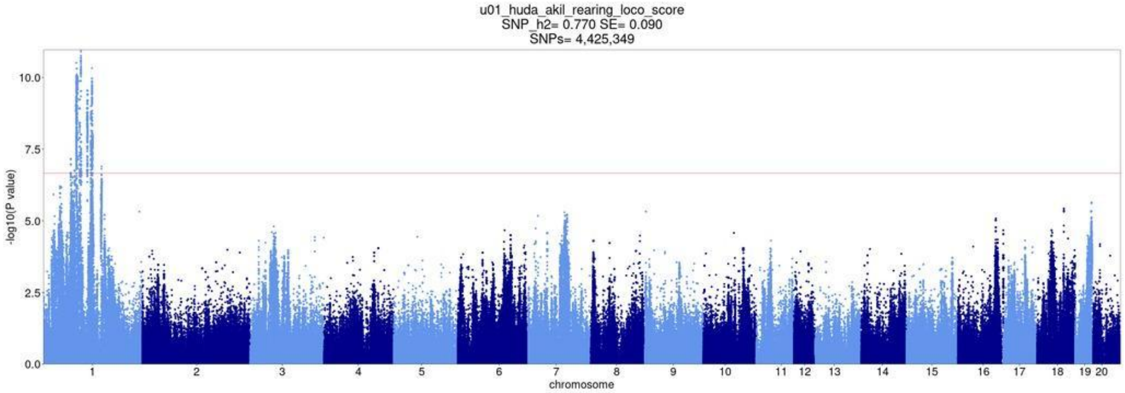


PavCA Response bias day6

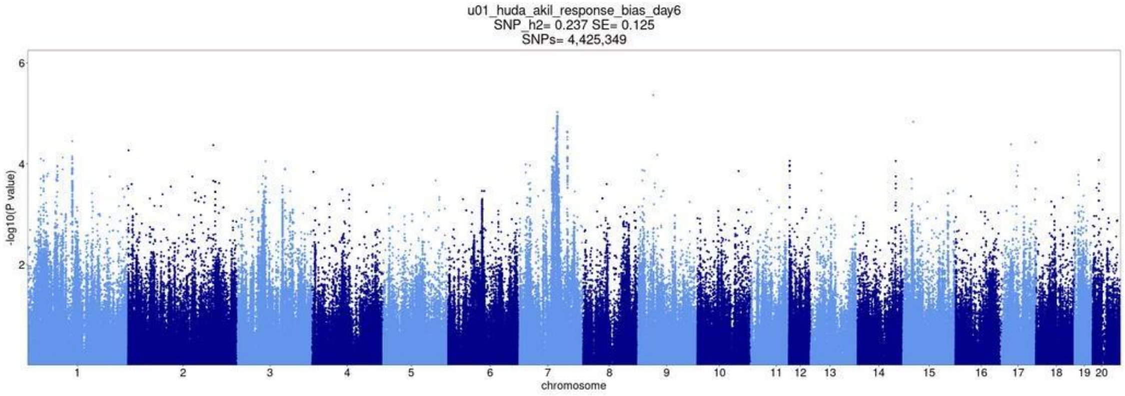

PavCA Response bias day7

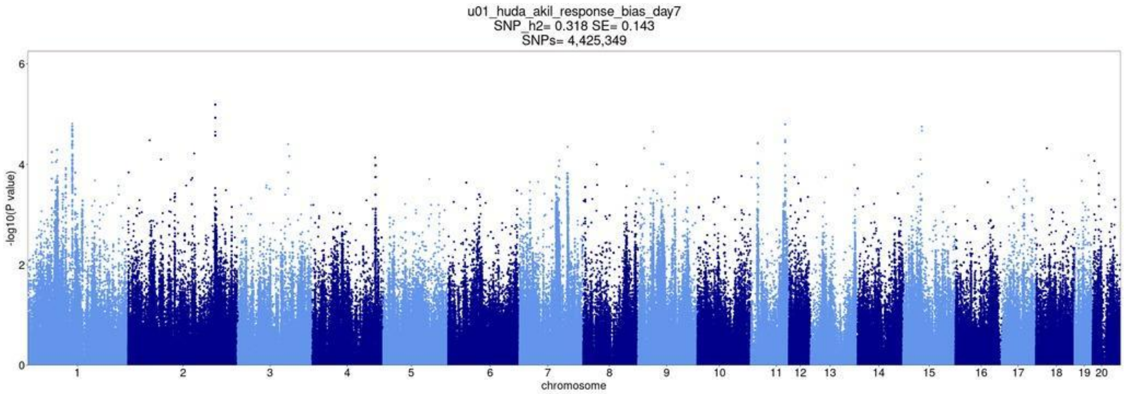


Total locomotor score

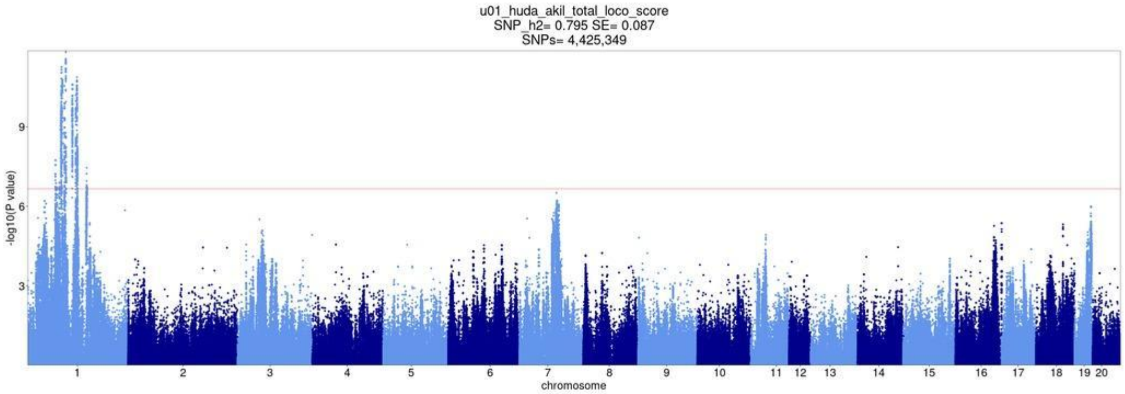

**Supplemental Figure 2.** Regional association plots for all QTLs.

EPM distance traveled: chr1 94,914,942

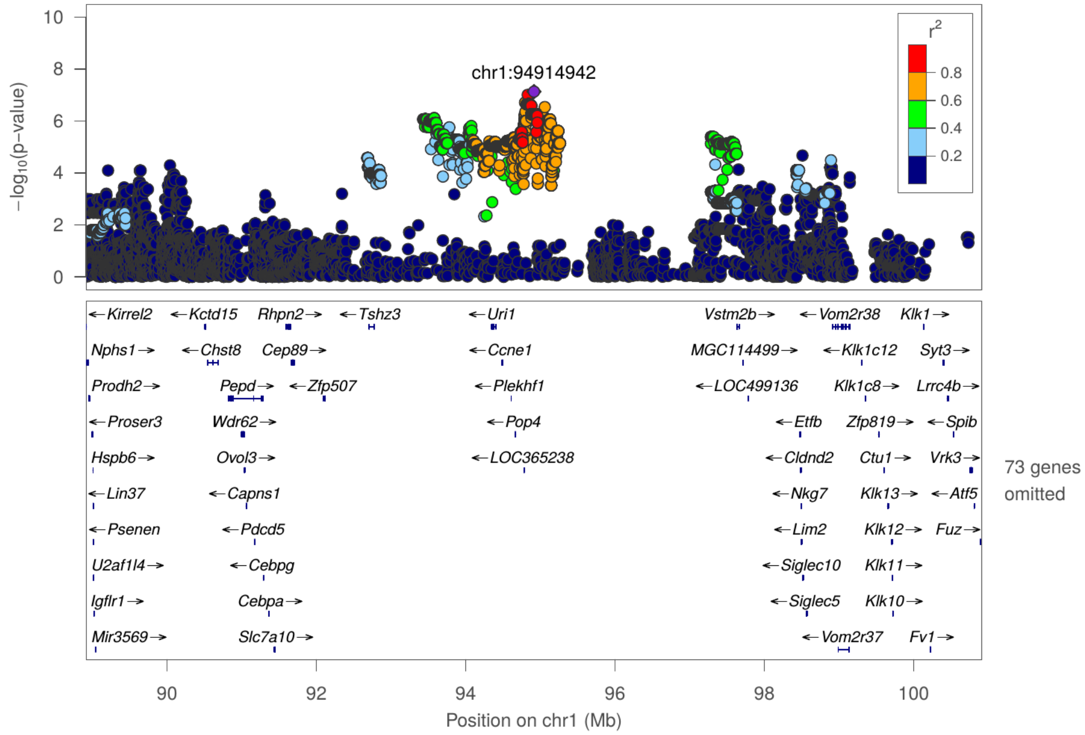

Lateral locomotor score: chr1 107,181,408

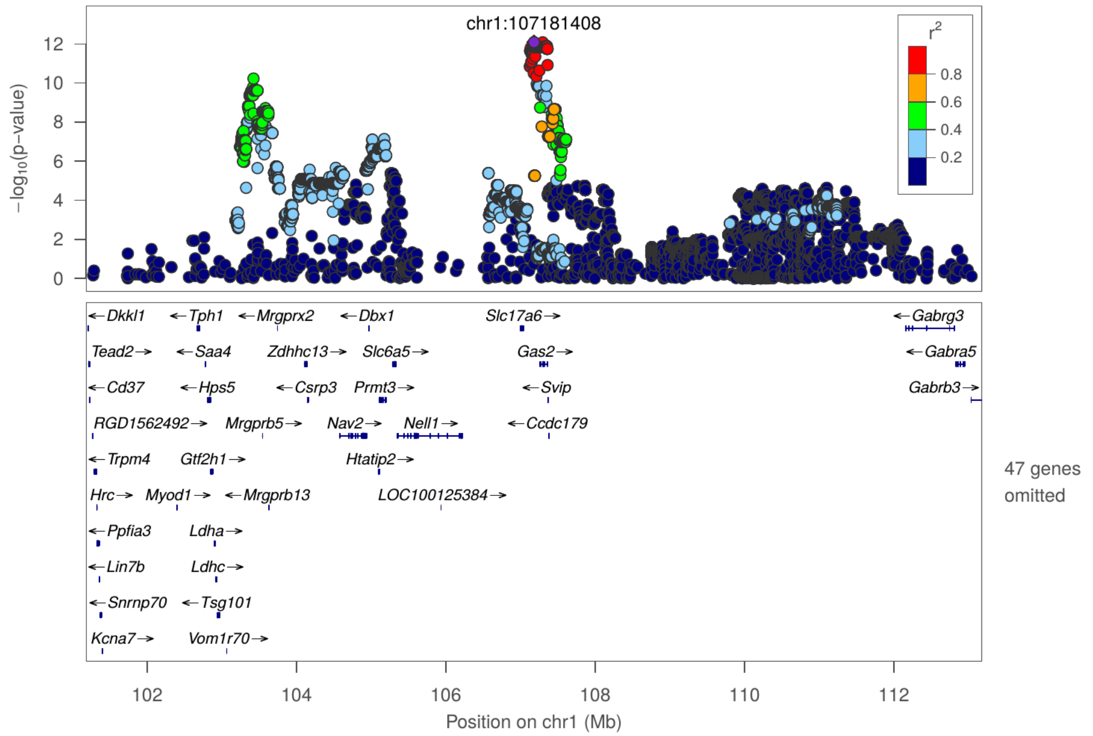

Lateral locomotor score: chr1 95,096,740

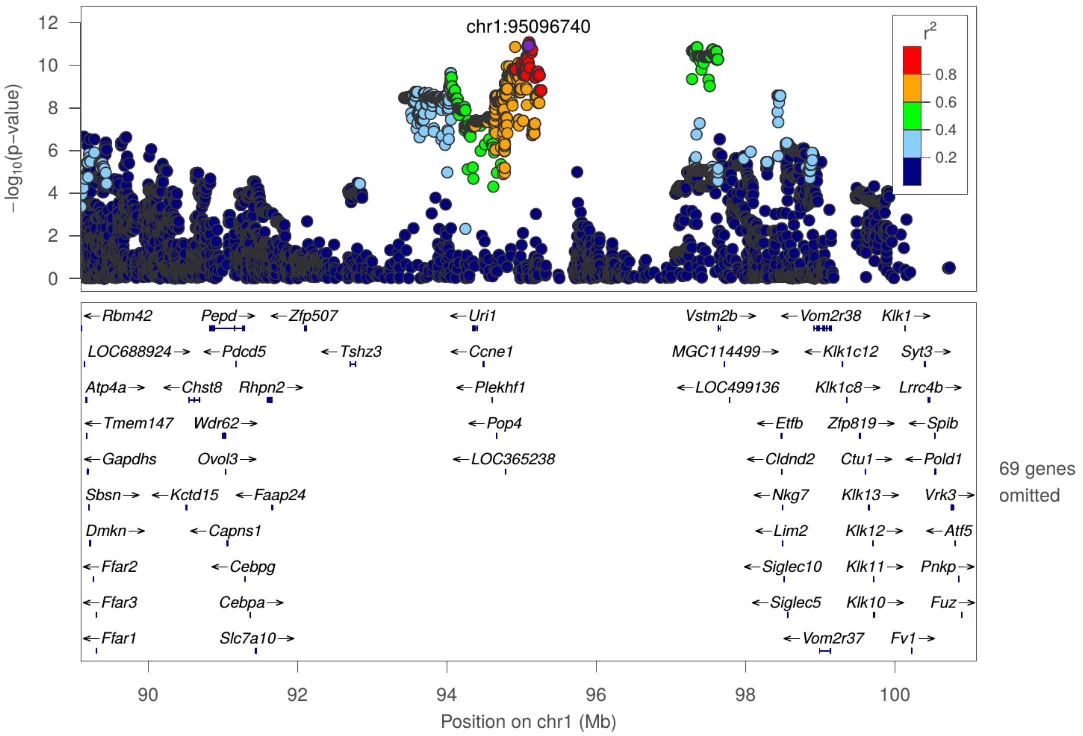

Lateral locomotor score: chr7 84,269,306

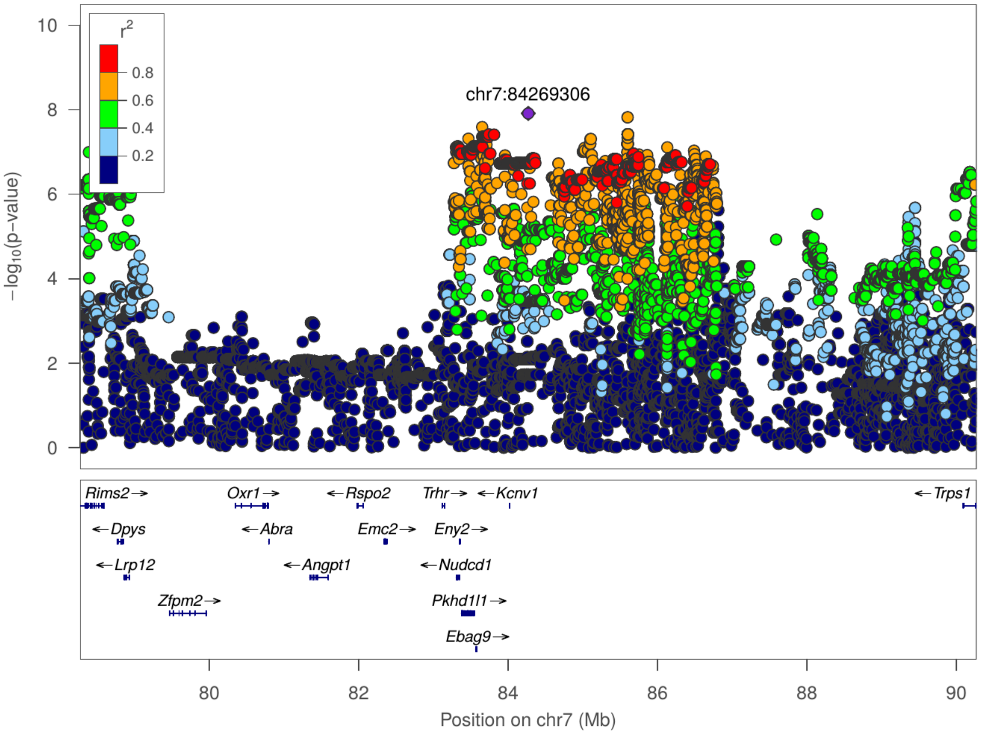

Open field boli: chr18 23,205,496

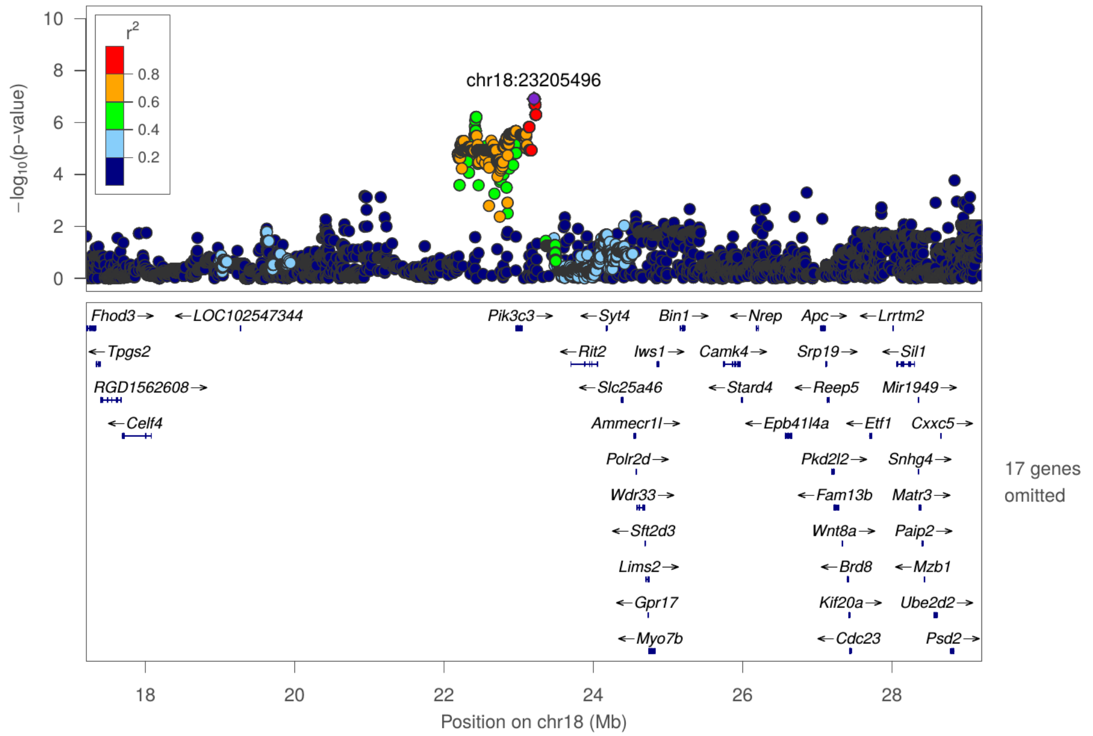

Open field distance traveled : chr1 135,389,579

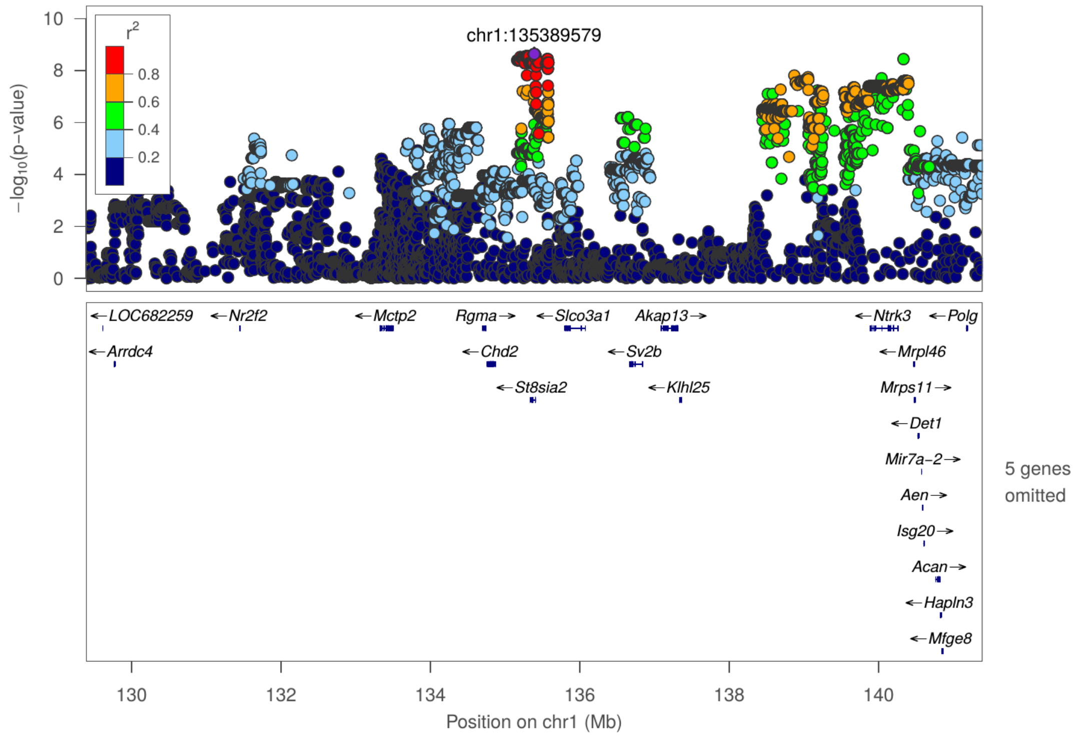

Rearing locomotor score: chr1 107,298,166

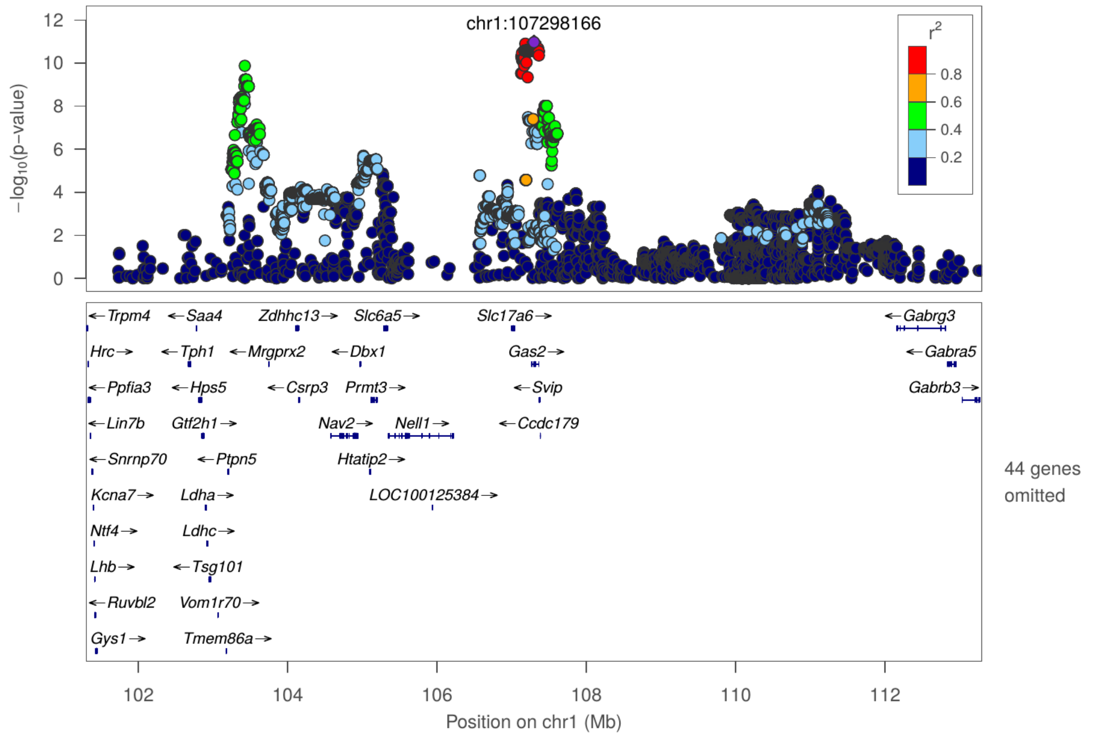

Rearing locomotor score: chr1 95,087,956

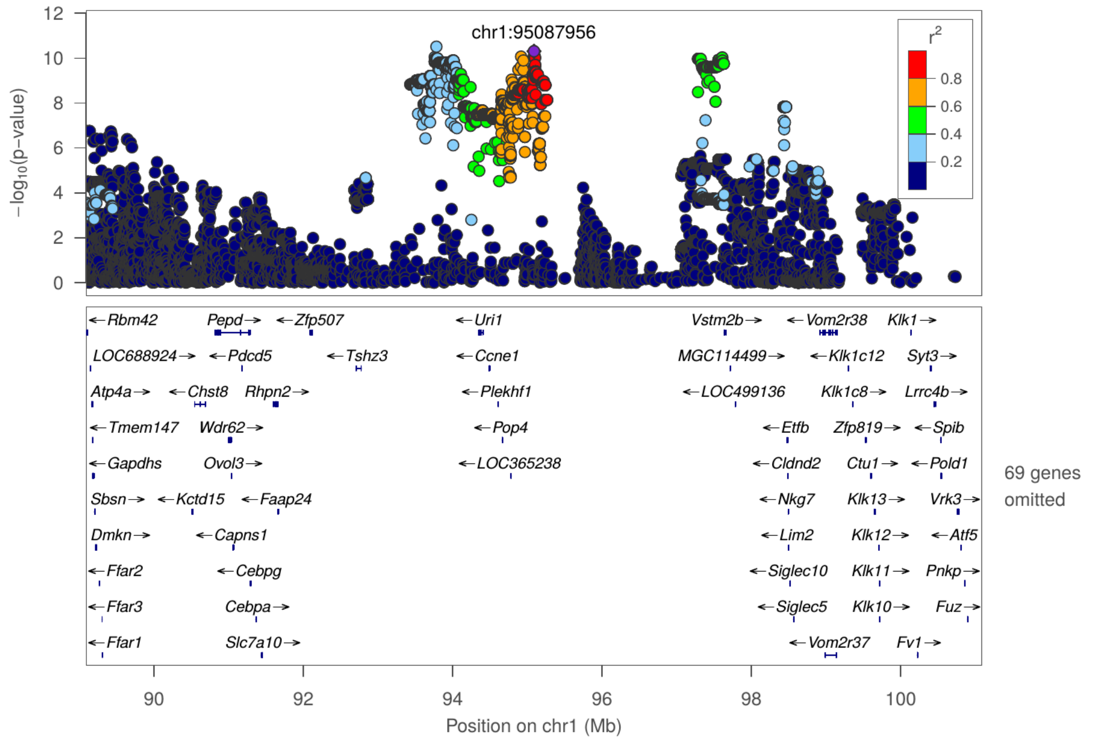

Total locomotor score: chr1 107,298,166

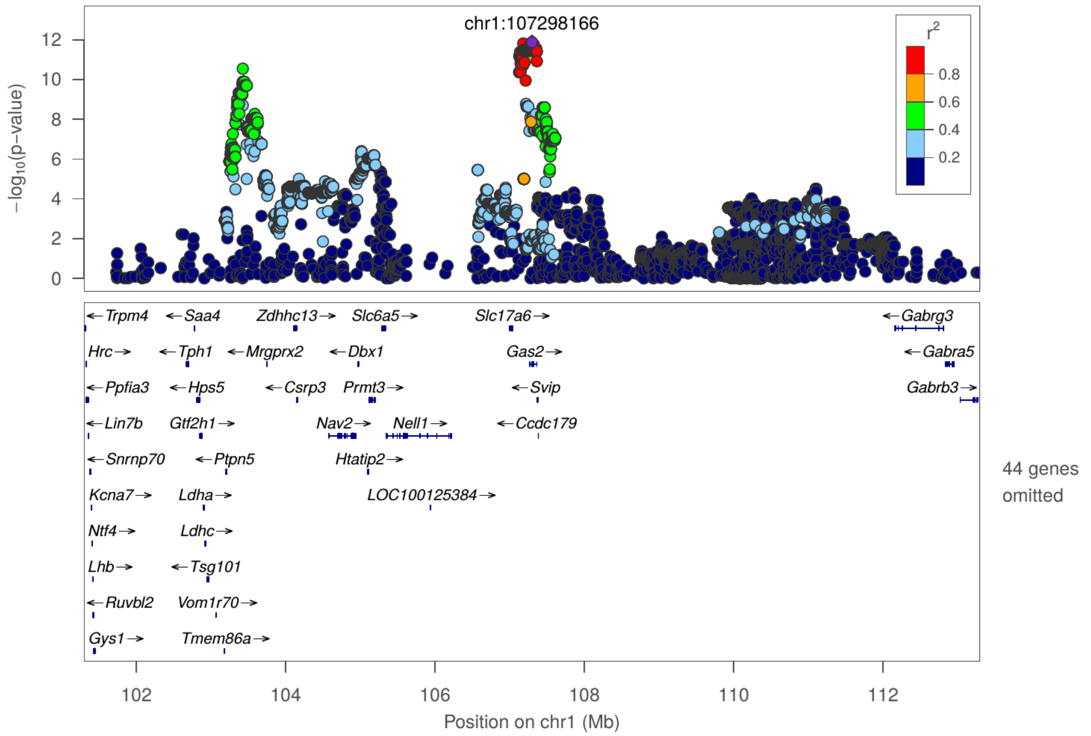

Total locomotor score: chr1 95,096,740

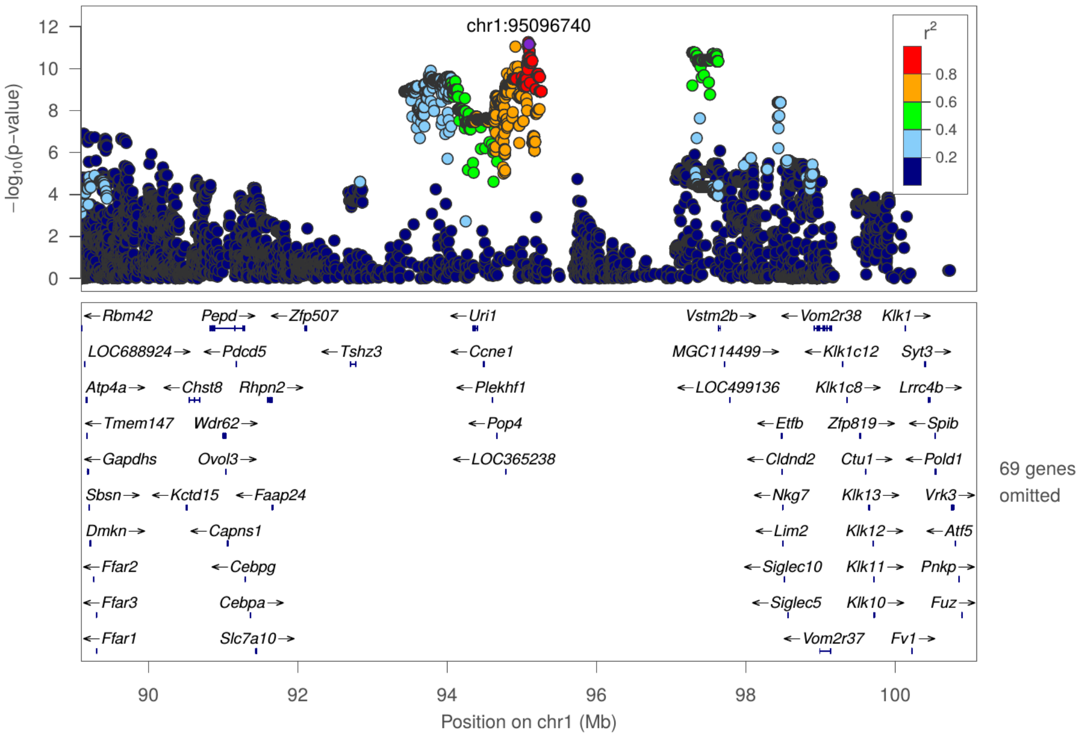

**Supplemental Table 1.**

Summary statistics for all behavioral traits.

| | A. Females | | | | | | B. Males | | | | | |
|---|---|---|---|---|---|---|---|---|---|---|---|---|
| **Traits** | **Stand. dev** | **Mean** | **n** | **Median** | **Minimum** | **Maximum** | **Stand. dev** | **Mean** | **n** | **Median** | **Minimum** | **Maximum** |
| Lateral locomotor score | 114.453 | 216.682 | 264 | 194 | 34 | 558 | 99.874 | 183.477 | 275 | 163.5 | 19 | 520 |
| Rearing locomotor score | 193.368 | 351.728 | 264 | 324 | 41 | 946 | 168.125 | 295.657 | 275 | 263 | 21 | 831 |
| Total locomotor score | 300.448 | 568.411 | 264 | 513 | 75 | 1461 | 259.304 | 479.134 | 275 | 432 | 40 | 1319 |
| EPM percent time in open arm | 10.881 | 16.759 | 264 | 15.67 | 0 | 45.03 | 10.125 | 11.075 | 275 | 8.3 | 0 | 46.15 |
| EPM boli | 0.648 | 0.161 | 264 | 0 | 0 | 5 | 2.035 | 1.251 | 275 | 0 | 0 | 8 |
| EPM distance traveled | 430.404 | 2202.693 | 264 | 2168.85 | 1221.67 | 3338.77 | 502.615 | 1943.112 | 275 | 1852.515 | 1103.76 | 5208.82 |
| EPM time immobile [s] | 18.486 | 86.597 | 264 | 86.29 | 40.74 | 147.15 | 25.146 | 113.833 | 275 | 113.76 | 63.86 | 179.08 |
| Open field Percent time in center | 2.391 | 3.145 | 264 | 2.467 | 0 | 12.467 | 2.759 | 2.91 | 275 | 2.067 | 0 | 14.933 |
| Open field boli | 1.433 | 1.018 | 264 | 0 | 0 | 6 | 1.861 | 1.922 | 275 | 2 | 0 | 6 |
| Open field distance traveled | 580.133 | 2635.145 | 264 | 2634.607 | 1307.886 | 4080.152 | 633.863 | 2400.69 | 275 | 2454.608 | 293.544 | 3587.956 |
| Open field time | 20.768 | 61.261 | 264 | 58.659 | 26.827 | 129.329 | 30.099 | 65.846 | 275 | 60.661 | 14.815 | 230.631 |

| | | | | | | | | | | | | |
|---|---|---|---|---|---|---|---|---|---|---|---|---|
| immobile [s] | | | | | | | | | | | | |
| PavCA index days 6and7 | 0.5 | 0.404 | 264 | 0.618 | -0.904 | 0.909 | 0.518 | -0.118 | 275 | -0.221 | -0.882 | 0.848 |
| PavCA Response bias day7 | 0.684 | 0.556 | 264 | 0.939 | -1 | 1 | 0.789 | -0.182 | 275 | -0.351 | -1 | 1 |
| PavCA prob diff day7 | 0.534 | 0.463 | 264 | 0.64 | -1 | 1 | 0.552 | -0.047 | 275 | -0.08 | -0.92 | 0.96 |
| PavCA latency score day7 | 0.341 | 0.263 | 264 | 0.306 | -0.715 | 0.756 | 0.355 | -0.043 | 275 | -0.07 | -0.758 | 0.7 |
| PavCA index day7only | 0.506 | 0.427 | 264 | 0.627 | -0.905 | 0.919 | 0.546 | -0.091 | 275 | -0.255 | -0.893 | 0.887 |
| PavCA Response bias day6 | 0.707 | 0.479 | 264 | 0.895 | -1 | 1 | 0.757 | -0.257 | 275 | -0.52 | -1 | 1 |
| PavCA prob diff day6 | 0.537 | 0.402 | 264 | 0.56 | -0.96 | 1 | 0.536 | -0.093 | 275 | -0.16 | -0.92 | 0.96 |
| PavCA latency score day6 | 0.355 | 0.202 | 264 | 0.244 | -0.765 | 0.852 | 0.337 | -0.083 | 275 | -0.116 | -0.748 | 0.602 |
| PavCA index day6only | 0.517 | 0.361 | 264 | 0.588 | -0.902 | 0.951 | 0.529 | -0.144 | 275 | -0.366 | -0.871 | 0.817 |

**Supplemental Table 2.**

Phenotypic and genetic correlation estimates between all traits.

| | | Phenotypic correlations | | Genetic correlations | | |
|---|---|---|---|---|---|---|
| **Trait 1** | **Trait 2** | **rho** | **Pval_pheno** | **rG** | **SE** | **Pval_gen** |
| **EPM boli** | **EPM dist.** | **-0.17** | **4.23E-04** | **-0.51** | **0.18** | **8.70E-03** |
| **EPM boli** | **EPM time imm.** | **0.15** | **3.13E-03** | **0.17** | **0.30** | **2.95E-01** |
| **EPM boli** | **PavCA Lat. D6** | **-0.06** | **3.75E-01** | **0.12** | **0.38** | **3.70E-01** |
| **EPM boli** | **PavCA Lat. D7** | **0.00** | **9.75E-01** | **0.32** | **0.33** | **1.68E-01** |
| **EPM boli** | **OF boli** | **NA** | **NA** | **0.45** | **0.29** | **5.77E-02** |
| **EPM boli** | **OF dist.** | **NA** | **NA** | **-0.36** | **0.26** | **7.95E-02** |
| **EPM boli** | **OF % time** | **NA** | **NA** | **0.04** | **0.45** | **4.63E-01** |
| **EPM boli** | **OF time imm.** | **NA** | **NA** | **0.38** | **0.29** | **8.23E-02** |
| **EPM boli** | **PavCA index D6** | **-0.08** | **2.33E-01** | **0.13** | **0.41** | **3.65E-01** |
| **EPM boli** | **PavCA index D7** | **-0.01** | **8.89E-01** | **0.24** | **0.34** | **2.35E-01** |
| **EPM boli** | **PavCA index D6D7** | **-0.04** | **5.97E-01** | **0.17** | **0.32** | **2.99E-01** |
| **EPM boli** | **PavCA ProbDiff D6** | **-0.06** | **3.83E-01** | **0.39** | **0.64** | **2.37E-01** |
| **EPM boli** | **PavCA ProbDiff D7** | **0.01** | **8.66E-01** | **0.42** | **0.39** | **1.38E-01** |
| **EPM boli** | **PavCA Resp. D6** | **-0.06** | **4.18E-01** | **0.37** | **0.34** | **1.34E-01** |
| **EPM boli** | **PavCA Resp. D7** | **0.00** | **9.83E-01** | **0.24** | **0.32** | **2.24E-01** |
| **EPM dist.** | **EPM time imm.** | **-0.77** | **0.00E+00** | **NA** | **NA** | **NA** |
| **EPM dist.** | **PavCA Lat. D6** | **0.26** | **1.51E-04** | **1.00** | **0.29** | **7.12E-05** |
| **EPM dist.** | **PavCA Lat. D7** | **0.20** | **3.71E-03** | **1.00** | **0.13** | **2.94E-05** |
| **EPM dist.** | **OF boli** | **NA** | **NA** | **-0.37** | **0.29** | **1.01E-01** |
| **EPM dist.** | **OF dist.** | **NA** | **NA** | **1.00** | **0.16** | **2.04E-05** |
| **EPM dist.** | **OF % time** | **NA** | **NA** | **0.64** | **0.45** | **9.34E-02** |
| **EPM dist.** | **OF time imm.** | **NA** | **NA** | **-0.83** | **0.20** | **4.29E-04** |

| | | | | | | |
|---|---|---|---|---|---|---|
| EPM dist. | PavCA index D6 | 0.26 | 1.53E-04 | 1.00 | 0.27 | 6.02E-05 |
| EPM dist. | PavCA index D7 | 0.22 | 1.21E-03 | 1.00 | 0.14 | 2.15E-05 |
| EPM dist. | PavCA index D6D7 | 0.25 | 2.26E-04 | 1.00 | 0.18 | 1.60E-05 |
| EPM dist. | PavCA ProbDiff D6 | 0.23 | 8.14E-04 | 1.00 | 0.31 | 5.41E-04 |
| EPM dist. | PavCA ProbDiff D7 | 0.21 | 1.94E-03 | 0.98 | 0.15 | 1.81E-04 |
| EPM dist. | PavCA Resp. D6 | 0.25 | 2.09E-04 | 0.90 | 0.14 | 1.39E-04 |
| EPM dist. | PavCA Resp. D7 | 0.28 | 4.24E-05 | 0.91 | 0.12 | 3.72E-05 |
| EPM % time | EPM boli | -0.07 | 1.79E-01 | -0.10 | 0.27 | 3.50E-01 |
| EPM % time | EPM dist. | 0.44 | 0.00E+00 | 0.71 | 0.16 | 1.37E-03 |
| EPM % time | EPM time imm. | -0.36 | 5.39E-14 | -0.64 | 0.24 | 3.34E-02 |
| EPM % time | PavCA Lat. D6 | 0.11 | 1.05E-01 | 0.86 | 0.37 | 1.52E-02 |
| EPM % time | PavCA Lat. D7 | 0.10 | 1.44E-01 | 0.78 | 0.26 | 7.55E-03 |
| EPM % time | OF boli | NA | NA | 0.35 | 0.38 | 1.72E-01 |
| EPM % time | OF dist. | NA | NA | 0.60 | 0.26 | 2.00E-02 |
| EPM % time | OF % time | NA | NA | 0.61 | 0.38 | 1.08E-01 |
| EPM % time | OF time imm. | NA | NA | -0.69 | 0.23 | 7.08E-03 |
| EPM % time | PavCA index D6 | 0.11 | 9.74E-02 | 0.98 | 0.41 | 9.15E-03 |
| EPM % time | PavCA index D7 | 0.10 | 1.57E-01 | 0.78 | 0.29 | 1.08E-02 |
| EPM % time | PavCA index D6D7 | 0.12 | 8.49E-02 | 0.82 | 0.27 | 6.58E-03 |
| EPM % time | PavCA ProbDiff D6 | 0.10 | 1.57E-01 | 1.00 | 0.59 | 1.62E-02 |
| EPM % time | PavCA ProbDiff D7 | 0.11 | 1.19E-01 | 0.81 | 0.29 | 1.04E-02 |
| EPM % time | PavCA Resp. D6 | 0.15 | 3.04E-02 | 0.55 | 0.30 | 4.77E-02 |
| EPM % time | PavCA Resp. D7 | 0.13 | 7.04E-02 | 0.65 | 0.26 | 1.85E-02 |
| EPM time imm. | PavCA Lat. D6 | -0.25 | 2.30E-04 | -1.00 | 0.21 | 7.79E-04 |

| | | | | | | |
|---|---|---|---|---|---|---|
| EPM time imm. | PavCA Lat. D7 | -0.21 | 2.86E-03 | -1.00 | 0.19 | 1.44E-04 |
| EPM time imm. | OF boli | NA | NA | 0.12 | 0.41 | 3.77E-01 |
| EPM time imm. | OF dist. | NA | NA | -1.00 | 0.28 | 6.09E-04 |
| EPM time imm. | OF % time | NA | NA | -1.00 | 0.62 | 4.26E-02 |
| EPM time imm. | OF time imm. | NA | NA | 1.00 | 0.25 | 6.28E-04 |
| EPM time imm. | PavCA index D6 | -0.25 | 2.32E-04 | -1.00 | 0.25 | 7.30E-04 |
| EPM time imm. | PavCA index D7 | -0.22 | 1.72E-03 | -1.00 | 0.19 | 2.02E-04 |
| EPM time imm. | PavCA index D6D7 | -0.25 | 3.42E-04 | -1.00 | 0.17 | 2.38E-04 |
| EPM time imm. | PavCA ProbDiff D6 | -0.25 | 2.83E-04 | -1.00 | 0.70 | 1.80E-02 |
| EPM time imm. | PavCA ProbDiff D7 | -0.23 | 6.98E-04 | -1.00 | 0.26 | 4.41E-04 |
| EPM time imm. | PavCA Resp. D6 | -0.25 | 2.64E-04 | -1.00 | 0.29 | 2.12E-04 |
| EPM time imm. | PavCA Resp. D7 | -0.26 | 1.73E-04 | -1.00 | 0.17 | 2.43E-05 |
| PavCA Lat. D6 | PavCA index D6 | 0.97 | 0.00E+00 | 0.98 | 0.02 | 0.00E+00 |
| PavCA Lat. D7 | PavCA Lat. D6 | 0.87 | 0.00E+00 | 1.00 | 0.05 | 1.15E-02 |
| PavCA Lat. D7 | PavCA index D6 | 0.86 | 0.00E+00 | 1.00 | 0.11 | 8.51E-02 |
| PavCA Lat. D7 | PavCA index D7 | 0.97 | 0.00E+00 | 1.00 | 0.01 | 0.00E+00 |

| | | | | | | |
|---|---|---|---|---|---|---|
| **PavCA Lat. D7** | **PavCA ProbDiff D6** | **0.86** | **0.00E+00** | **1.00** | **0.49** | **1.11E-01** |
| **PavCA Lat. D7** | **PavCA Resp. D6** | **0.71** | **0.00E+00** | **0.93** | **0.11** | **8.18E-03** |
| **Lat. Loco. Score** | **EPM boli** | **-0.10** | **3.97E-02** | **-0.15** | **0.19** | **2.13E-01** |
| **Lat. Loco. Score** | **EPM dist.** | **0.40** | **0.00E+00** | **0.76** | **0.10** | **8.23E-07** |
| **Lat. Loco. Score** | **EPM % time** | **0.24** | **5.39E-07** | **0.70** | **0.16** | **4.04E-04** |
| **Lat. Loco. Score** | **EPM time imm.** | **-0.36** | **7.80E-14** | **-1.00** | **0.13** | **9.67E-07** |
| **Lat. Loco. Score** | **PavCA Lat. D6** | **0.41** | **1.19E-09** | **0.90** | **0.27** | **2.18E-03** |
| **Lat. Loco. Score** | **PavCA Lat. D7** | **0.38** | **1.46E-08** | **0.95** | **0.18** | **9.04E-05** |
| **Lat. Loco. Score** | **OF boli** | **NA** | **NA** | **-0.12** | **0.23** | **2.85E-01** |
| **Lat. Loco. Score** | **OF dist.** | **NA** | **NA** | **0.88** | **0.13** | **2.12E-07** |
| **Lat. Loco. Score** | **OF % time** | **NA** | **NA** | **0.41** | **0.36** | **1.19E-01** |
| **Lat. Loco. Score** | **OF time imm.** | **NA** | **NA** | **-0.62** | **0.19** | **9.17E-04** |
| **Lat. Loco. Score** | **PavCA index D6** | **0.43** | **1.23E-10** | **1.00** | **0.31** | **4.73E-04** |
| **Lat. Loco. Score** | **PavCA index D7** | **0.39** | **8.68E-09** | **0.95** | **0.20** | **1.79E-04** |
| **Lat. Loco. Score** | **PavCA index D6D7** | **0.43** | **2.01E-10** | **0.88** | **0.18** | **3.01E-04** |
| **Lat. Loco. Score** | **PavCA ProbDiff D6** | **0.42** | **4.15E-10** | **1.00** | **0.54** | **1.15E-03** |

| | | | | | | |
|---|---|---|---|---|---|---|
| Lat. Loco. Score | PavCA ProbDiff D7 | 0.40 | 5.20E-09 | 1.00 | 0.28 | 6.82E-05 |
| Lat. Loco. Score | Rear. Loco. Score | 0.88 | 0.00E+00 | 0.97 | 0.01 | 0.00E+00 |
| Lat. Loco. Score | PavCA Resp. D6 | 0.48 | 0.00E+00 | 1.00 | 0.13 | 7.22E-07 |
| Lat. Loco. Score | PavCA Resp. D7 | 0.43 | 1.35E-10 | 1.00 | 0.16 | 3.04E-05 |
| Lat. Loco. Score | Tot. Loco. Score | 0.94 | 0.00E+00 | 0.99 | 0.01 | 0.00E+00 |
| OF boli | PavCA Lat. D6 | NA | NA | -0.29 | 0.46 | 2.64E-01 |
| OF boli | PavCA Lat. D7 | NA | NA | 0.09 | 0.42 | 4.13E-01 |
| OF boli | OF dist. | -0.20 | 3.68E-03 | -0.35 | 0.22 | 6.17E-02 |
| OF boli | OF time imm. | 0.03 | 7.08E-01 | 0.03 | 0.26 | 4.51E-01 |
| OF boli | PavCA index D6 | NA | NA | -0.06 | 0.51 | 4.51E-01 |
| OF boli | PavCA index D7 | NA | NA | 0.14 | 0.44 | 3.70E-01 |
| OF boli | PavCA index D6D7 | NA | NA | 0.02 | 0.41 | 4.84E-01 |
| OF boli | PavCA ProbDiff D6 | NA | NA | -0.10 | 0.67 | 4.42E-01 |
| OF boli | PavCA ProbDiff D7 | NA | NA | 0.30 | 0.49 | 2.67E-01 |
| OF boli | PavCA Resp. D6 | NA | NA | -0.11 | 0.40 | 3.82E-01 |
| OF boli | PavCA Resp. D7 | NA | NA | 0.27 | 0.38 | 2.31E-01 |
| OF dist. | PavCA Lat. D6 | NA | NA | 0.99 | 0.33 | 5.99E-03 |
| OF dist. | PavCA Lat. D7 | NA | NA | 0.91 | 0.25 | 2.91E-03 |
| OF dist. | OF time imm. | -0.61 | 1.72E-23 | -0.86 | 0.08 | 3.06E-06 |
| OF dist. | PavCA index D6 | NA | NA | 0.96 | 0.34 | 8.57E-03 |
| OF dist. | PavCA index D7 | NA | NA | 0.93 | 0.26 | 3.52E-03 |
| OF dist. | PavCA index D6D7 | NA | NA | 0.89 | 0.26 | 3.54E-03 |

| | | | | | | |
|---|---|---|---|---|---|---|
| OF dist. | PavCA ProbDiff D6 | NA | NA | 1.00 | 0.47 | 1.63E-02 |
| OF dist. | PavCA ProbDiff D7 | NA | NA | 0.96 | 0.28 | 4.78E-03 |
| OF dist. | PavCA Resp. D6 | NA | NA | 0.90 | 0.30 | 4.82E-03 |
| OF dist. | PavCA Resp. D7 | NA | NA | 0.79 | 0.27 | 6.28E-03 |
| OF % time | PavCA Lat. D6 | NA | NA | 0.19 | 0.80 | 4.29E-01 |
| OF % time | PavCA Lat. D7 | NA | NA | -0.37 | 0.76 | 3.52E-01 |
| OF % time | OF boli | -0.09 | 2.01E-01 | -0.37 | 0.43 | 1.82E-01 |
| OF % time | OF dist. | 0.41 | 2.56E-10 | 0.74 | 0.31 | 3.49E-02 |
| OF % time | OF time imm. | -0.30 | 5.05E-06 | -0.44 | 0.33 | 1.23E-01 |
| OF % time | PavCA index D6 | NA | NA | 0.12 | 0.86 | 4.60E-01 |
| OF % time | PavCA index D7 | NA | NA | -0.26 | 0.79 | 3.98E-01 |
| OF % time | PavCA index D6D7 | NA | NA | 0.17 | 0.67 | 4.20E-01 |
| OF % time | PavCA ProbDiff D6 | NA | NA | -0.01 | 1.21 | 4.97E-01 |
| OF % time | PavCA ProbDiff D7 | NA | NA | -0.74 | 0.86 | 2.59E-01 |
| OF % time | PavCA Resp. D6 | NA | NA | 0.77 | 0.62 | 1.41E-01 |
| OF % time | PavCA Resp. D7 | NA | NA | 0.53 | 0.50 | 1.94E-01 |
| OF time imm. | PavCA Lat. D6 | NA | NA | -0.49 | 0.43 | 1.35E-01 |
| OF time imm. | PavCA Lat. D7 | NA | NA | -0.40 | 0.36 | 1.51E-01 |
| OF time imm. | PavCA index D6 | NA | NA | -0.40 | 0.45 | 2.00E-01 |
| OF time imm. | PavCA index D7 | NA | NA | -0.41 | 0.37 | 1.53E-01 |
| OF time imm. | PavCA index D6D7 | NA | NA | -0.43 | 0.36 | 1.26E-01 |

| | | | | | | |
|---|---|---|---|---|---|---|
| OF time imm. | PavCA ProbDiff D6 | NA | NA | -0.38 | 0.60 | 2.73E-01 |
| OF time imm. | PavCA ProbDiff D7 | NA | NA | -0.40 | 0.41 | 1.82E-01 |
| OF time imm. | PavCA Resp. D6 | NA | NA | -0.40 | 0.37 | 1.35E-01 |
| OF time imm. | PavCA Resp. D7 | NA | NA | -0.44 | 0.32 | 9.64E-02 |
| PavCA index D7 | PavCA Lat. D6 | 0.85 | 0.00E+00 | 1.00 | 0.07 | 2.36E-02 |
| PavCA index D7 | PavCA index D6 | 0.87 | 0.00E+00 | NA | NA | NA |
| PavCA index D7 | PavCA ProbDiff D6 | 0.86 | 0.00E+00 | 1.00 | 0.47 | 4.07E-01 |
| PavCA index D7 | PavCA Resp. D6 | 0.75 | 0.00E+00 | 0.91 | 0.12 | 1.55E-02 |
| PavCA index D6D7 | PavCA Lat. D6 | 0.94 | 0.00E+00 | NA | NA | NA |
| PavCA index D6D7 | PavCA Lat. D7 | 0.94 | 0.00E+00 | NA | NA | NA |
| PavCA index D6D7 | PavCA index D6 | 0.97 | 0.00E+00 | NA | NA | NA |
| PavCA index D6D7 | PavCA index D7 | 0.96 | 0.00E+00 | NA | NA | NA |
| PavCA index D6D7 | PavCA ProbDiff D6 | 0.95 | 0.00E+00 | 1.00 | 0.07 | 0.00E+00 |
| PavCA index D6D7 | PavCA ProbDiff D7 | 0.94 | 0.00E+00 | NA | NA | NA |
| PavCA index D6D7 | PavCA Resp. D6 | 0.84 | 0.00E+00 | 0.83 | 0.14 | 3.25E-02 |
| PavCA index D6D7 | PavCA Resp. D7 | 0.83 | 0.00E+00 | 0.92 | 0.09 | 9.06E-03 |

| | | | | | | |
|---|---|---|---|---|---|---|
| PavCA ProbDiff D6 | PavCA Lat. D6 | 0.97 | 0.00E+00 | 0.98 | 0.05 | 0.00E+00 |
| PavCA ProbDiff D6 | PavCA index D6 | 0.99 | 0.00E+00 | 1.00 | 0.04 | 0.00E+00 |
| PavCA ProbDiff D7 | PavCA Lat. D6 | 0.83 | 0.00E+00 | 0.98 | 0.06 | 4.89E-02 |
| PavCA ProbDiff D7 | PavCA Lat. D7 | 0.96 | 0.00E+00 | NA | NA | NA |
| PavCA ProbDiff D7 | PavCA index D6 | 0.85 | 0.00E+00 | 1.00 | 0.07 | 4.07E-02 |
| PavCA ProbDiff D7 | PavCA index D7 | 0.98 | 0.00E+00 | 0.98 | 0.02 | 0.00E+00 |
| PavCA ProbDiff D7 | PavCA ProbDiff D6 | 0.85 | 0.00E+00 | NA | NA | NA |
| PavCA ProbDiff D7 | PavCA Resp. D6 | 0.71 | 0.00E+00 | 1.00 | 0.13 | 6.24E-03 |
| Rear. Loco. Score | EPM boli | -0.14 | 4.06E-03 | -0.16 | 0.19 | 2.02E-01 |
| Rear. Loco. Score | EPM dist. | 0.41 | 0.00E+00 | 0.73 | 0.10 | 3.47E-06 |
| Rear. Loco. Score | EPM % time | 0.25 | 1.70E-07 | 0.64 | 0.17 | 9.78E-04 |
| Rear. Loco. Score | EPM time imm. | -0.43 | 0.00E+00 | -1.00 | 0.14 | 2.03E-06 |
| Rear. Loco. Score | PavCA Lat. D6 | 0.41 | 6.35E-10 | 0.92 | 0.23 | 4.91E-04 |
| Rear. Loco. Score | PavCA Lat. D7 | 0.39 | 7.49E-09 | 0.90 | 0.17 | 8.03E-05 |
| Rear. Loco. Score | OF boli | NA | NA | -0.19 | 0.23 | 2.00E-01 |
| Rear. Loco. Score | OF dist. | NA | NA | 0.83 | 0.14 | 1.41E-06 |

| | | | | | | |
|---|---|---|---|---|---|---|
| Rear. Loco. Score | OF % time | NA | NA | 0.69 | 0.36 | 2.32E-02 |
| Rear. Loco. Score | OF time imm. | NA | NA | -0.58 | 0.20 | 2.79E-03 |
| Rear. Loco. Score | PavCA index D6 | 0.43 | 1.39E-10 | 1.00 | 0.28 | 1.95E-04 |
| Rear. Loco. Score | PavCA index D7 | 0.39 | 1.12E-08 | 0.90 | 0.19 | 2.15E-04 |
| Rear. Loco. Score | PavCA index D6D7 | 0.43 | 2.43E-10 | 0.85 | 0.18 | 2.39E-04 |
| Rear. Loco. Score | PavCA ProbDiff D6 | 0.41 | 6.06E-10 | 1.00 | 0.53 | 4.77E-04 |
| Rear. Loco. Score | PavCA ProbDiff D7 | 0.38 | 1.38E-08 | 1.00 | 0.22 | 1.41E-04 |
| Rear. Loco. Score | PavCA Resp. D6 | 0.47 | 0.00E+00 | 1.00 | 0.12 | 6.06E-07 |
| Rear. Loco. Score | PavCA Resp. D7 | 0.41 | 1.29E-09 | 0.91 | 0.17 | 1.77E-04 |
| Rear. Loco. Score | Tot. Loco. Score | 0.98 | 0.00E+00 | 1.00 | 0.00 | 0.00E+00 |
| PavCA Resp. D6 | PavCA Lat. D6 | 0.79 | 0.00E+00 | 0.74 | 0.21 | 8.58E-02 |
| PavCA Resp. D6 | PavCA index D6 | 0.86 | 0.00E+00 | 0.79 | 0.19 | 8.30E-02 |
| PavCA Resp. D6 | PavCA ProbDiff D6 | 0.82 | 0.00E+00 | 1.00 | 0.37 | 3.79E-02 |
| PavCA Resp. D7 | PavCA Lat. D6 | 0.68 | 0.00E+00 | 0.89 | 0.17 | 2.91E-02 |
| PavCA Resp. D7 | PavCA Lat. D7 | 0.77 | 0.00E+00 | 0.99 | 0.08 | 3.55E-03 |
| PavCA Resp. D7 | PavCA index D6 | 0.74 | 0.00E+00 | 1.00 | 0.15 | 1.29E-02 |

| | | | | | | |
|---|---|---|---|---|---|---|
| PavCA Resp. D7 | PavCA index D7 | 0.85 | 0.00E+00 | 0.98 | 0.06 | 4.97E-03 |
| PavCA Resp. D7 | PavCA ProbDiff D6 | 0.71 | 0.00E+00 | 1.00 | 0.61 | 3.81E-02 |
| PavCA Resp. D7 | PavCA ProbDiff D7 | 0.81 | 0.00E+00 | 1.00 | 0.19 | 8.69E-03 |
| PavCA Resp. D7 | PavCA Resp. D6 | 0.80 | 0.00E+00 | 1.00 | 0.07 | 1.64E-03 |
| Tot. Loco. Score | EPM boli | -0.13 | 7.56E-03 | -0.17 | 0.19 | 1.90E-01 |
| Tot. Loco. Score | EPM dist. | 0.42 | 0.00E+00 | 0.74 | 0.10 | 1.30E-06 |
| Tot. Loco. Score | EPM % time | 0.26 | 7.22E-08 | 0.67 | 0.16 | 5.28E-04 |
| Tot. Loco. Score | EPM time imm. | -0.42 | 0.00E+00 | -1.00 | 0.14 | 9.84E-07 |
| Tot. Loco. Score | PavCA Lat. D6 | 0.42 | 2.03E-10 | 0.93 | 0.24 | 6.55E-04 |
| Tot. Loco. Score | PavCA Lat. D7 | 0.40 | 3.25E-09 | 0.93 | 0.17 | 6.60E-05 |
| Tot. Loco. Score | OF boli | NA | NA | -0.17 | 0.23 | 2.24E-01 |
| Tot. Loco. Score | OF dist. | NA | NA | 0.86 | 0.13 | 4.02E-07 |
| Tot. Loco. Score | OF % time | NA | NA | 0.59 | 0.36 | 4.48E-02 |
| Tot. Loco. Score | OF time imm. | NA | NA | -0.60 | 0.19 | 1.57E-03 |
| Tot. Loco. Score | PavCA index D6 | 0.44 | 2.55E-11 | 1.00 | 0.28 | 1.85E-04 |
| Tot. Loco. Score | PavCA index D7 | 0.40 | 3.42E-09 | 0.93 | 0.19 | 1.58E-04 |

| | | | | | | |
|---|---|---|---|---|---|---|
| Tot. Loco. Score | PavCA index D6D7 | 0.44 | 5.24E-11 | 0.87 | 0.18 | 1.91E-04 |
| Tot. Loco. Score | PavCA ProbDiff D6 | 0.43 | 1.36E-10 | 1.00 | 0.53 | 4.49E-04 |
| Tot. Loco. Score | PavCA ProbDiff D7 | 0.40 | 3.35E-09 | 1.00 | 0.24 | 7.97E-05 |
| Tot. Loco. Score | PavCA Resp. D6 | 0.49 | 0.00E+00 | 1.00 | 0.12 | 2.45E-07 |
| Tot. Loco. Score | PavCA Resp. D7 | 0.43 | 1.10E-10 | 0.97 | 0.16 | 7.24E-05 |
| Lat. Loco. Score | Lat. Loco. Score | 1.00 | 0.00E+00 | 1.00 | NA | 0.00E+00 |
| Rear. Loco. Score | Rear. Loco. Score | 1.00 | 0.00E+00 | 1.00 | NA | 0.00E+00 |
| Tot. Loco. Score | Tot. Loco. Score | 1.00 | 0.00E+00 | 1.00 | NA | 0.00E+00 |
| PavCA Resp. D6 | PavCA Resp. D6 | 1.00 | 0.00E+00 | 1.00 | NA | 0.00E+00 |
| PavCA Resp. D7 | PavCA Resp. D7 | 1.00 | 0.00E+00 | 1.00 | NA | 0.00E+00 |
| PavCA ProbDiff D6 | PavCA ProbDiff D6 | 1.00 | 0.00E+00 | 1.00 | NA | 0.00E+00 |
| PavCA ProbDiff D7 | PavCA ProbDiff D7 | 1.00 | 0.00E+00 | 1.00 | NA | 0.00E+00 |
| PavCA Lat. D6 | PavCA Lat. D6 | 1.00 | 0.00E+00 | 1.00 | NA | 0.00E+00 |
| PavCA Lat. D7 | PavCA Lat. D7 | 1.00 | 0.00E+00 | 1.00 | NA | 0.00E+00 |
| PavCA index D6 | PavCA index D6 | 1.00 | 0.00E+00 | 1.00 | NA | 0.00E+00 |
| PavCA index D7 | PavCA index D7 | 1.00 | 0.00E+00 | 1.00 | NA | 0.00E+00 |

| | | | | | | |
|---|---|---|---|---|---|---|
| PavCA index D6D7 | PavCA index D6D7 | 1.00 | 0.00E+00 | 1.00 | NA | 0.00E+00 |
| EPM % time | EPM % time | 1.00 | 0.00E+00 | 1.00 | NA | 0.00E+00 |
| EPM boli | EPM boli | 1.00 | 0.00E+00 | 1.00 | NA | 0.00E+00 |
| EPM dist. | EPM dist. | 1.00 | 0.00E+00 | 1.00 | NA | 0.00E+00 |
| EPM time imm. | EPM time imm. | 1.00 | 0.00E+00 | 1.00 | NA | 0.00E+00 |
| OF % time | OF % time | 1.00 | 0.00E+00 | 1.00 | NA | 0.00E+00 |
| OF boli | OF boli | 1.00 | 0.00E+00 | 1.00 | NA | 0.00E+00 |
| OF dist. | OF dist. | 1.00 | 0.00E+00 | 1.00 | NA | 0.00E+00 |
| OF time imm. | OF time imm. | 1.00 | 0.00E+00 | 1.00 | NA | 0.00E+00 |

**Supplemental Table 3.** Genotypes of the $F_0$ rats at moderate impact variants

| | | | | | | | | | | | | | | | | | | | | | | |
|---|---|---|---|---|---|---|---|---|---|---|---|---|---|---|---|---|---|---|---|---|---|---|
| | | | bLR-F | bHR-M | bHR-F | bLR-M | bLR-F | bLR-M | bHR-M | bHR-F | bHR-M | bHR-F | bLR-F | bHR-M | bHR-F | bLR-M | bHR-M | bHR-F | bLR-F | bLR-M | bLR-F | bLR-M |
| Trait | QTL | Candidate gene | 5739-JL-0021 | 5739-JL-0022 | 5739-JL-0023 | 5739-JL-0024 | 5739-JL-0025 | 5739-JL-0026 | 5739-JL-0027 | 5739-JL-0028 | 5739-JL-0029 | 5739-JL-0030 | 5739-JL-0031 | 5739-JL-0032 | 5739-JL-0033 | 5739-JL-0034 | 5739-JL-0035 | 5739-JL-0036 | 5739-JL-0037 | 5739-JL-0038 | 5739-JL-0039 | 5739-JL-0040 |
| EPM distance traveled | chr1:94 Mb | *Plekhf1* | 1/1 | 0/0 | 0/1 | 1/1 | 1/1 | 1/1 | 0/0 | 0/0 | 0/0 | 0/1 | 1/1 | 0/1 | 0/0 | 1/1 | 0/1 | 0/1 | 0/1 | 0/1 | 0/1 | 1/1 |
| Lateral locomotor score | chr1:95 Mb | *Plekhf1* | 1/1 | 0/0 | 0/1 | 1/1 | 1/1 | 1/1 | 0/0 | 0/0 | 0/0 | 0/1 | 1/1 | 0/1 | 0/0 | 1/1 | 0/1 | 0/1 | 0/1 | 0/1 | 0/1 | 1/1 |
| Lateral locomotor score | chr7:84 Mb | *Pkhd1l1* | 0/0 | 1/1 | 1/1 | 0/0 | 0/0 | 0/0 | 1/1 | 1/1 | 1/1 | 1/1 | 0/0 | 1/1 | 1/1 | 0/0 | 1/1 | 1/1 | 0/0 | 0/0 | 0/0 | 0/0 |
| Rearing locomoto | chr1:95 Mb | *Plekhf1* | 1/1 | 0/0 | 0/1 | 1/1 | 1/1 | 1/1 | 0/0 | 0/0 | 0/0 | 0/1 | 1/1 | 0/1 | 0/0 | 1/1 | 0/1 | 0/1 | 0/1 | 0/1 | 0/1 | 1/1 |

| | | | | | | | | | | | | | | | | | | | | | | |
|---|---|---|---|---|---|---|---|---|---|---|---|---|---|---|---|---|---|---|---|---|---|---|
| r score | | | | | | | | | | | | | | | | | | | | | | |
| Total locomotor score | chr1: 95 Mb | *Plekhf1* | 1/1 | 0/0 | 0/1 | 1/1 | 1/1 | 1/1 | 0/0 | 0/0 | 0/0 | 0/1 | 1/1 | 0/1 | 0/0 | 1/1 | 0/1 | 0/1 | 0/1 | 0/1 | 0/1 | 1/1 |

**Supplemental Table 4.** Genotypes of the $F_0$ rats at QTL peak markers.

| | | | | **bLR-F** | **bHR-M** | **bHR-F** | **bLR-M** | **bLR-F** | **bLR-M** | **bHR-M** | **bHR-F** | **bHR-M** | **bHR-F** | **bLR-F** | **bHR-M** | **bHR-F** | **bLR-M** | **bHR-M** | **bHR-F** | **bLR-F** | **bLR-M** | **bLR-F** | **bLR-M** |
|---|---|---|---|---|---|---|---|---|---|---|---|---|---|---|---|---|---|---|---|---|---|---|---|
| | | **REF** | **ALT** | **5739-JL-0021** | **5739-JL-0022** | **5739-JL-0023** | **5739-JL-0024** | **5739-JL-0025** | **5739-JL-0026** | **5739-JL-0027** | **5739-JL-0028** | **5739-JL-0029** | **5739-JL-0030** | **5739-JL-0031** | **5739-JL-0032** | **5739-JL-0033** | **5739-JL-0034** | **5739-JL-0035** | **5739-JL-0036** | **5739-JL-0037** | **5739-JL-0038** | **5739-JL-0039** | **5739-JL-0040** |
| **Trait** | **topSNP** | T | C | 1/1 | 0/0 | 0/1 | 1/1 | 1/1 | 1/1 | 0/0 | 0/0 | 0/0 | 0/1 | 1/1 | 0/1 | 0/1 | 1/1 | 0/1 | 0/1 | 0/1 | 0/1 | 1/1 | 1/1 |
| EPM distance traveled | chr1:94,914,942 | T | C | 0/0 | 0/0 | 0/0 | 0/0 | 0/0 | 0/0 | 0/0 | 0/1 | 0/0 | 0/1 | 0/0 | 0/0 | 0/0 | 0/0 | 0/0 | 0/1 | 0/0 | 0/0 | 0/0 | 0/0 |
| Lateral locomotor score | chr1:107,181,408 | T | G | 1/1 | 0/0 | 0/0 | 1/1 | 1/1 | 1/1 | 0/0 | 0/0 | 0/0 | 0/0 | 1/1 | 0/1 | 0/1 | 1/1 | 0/0 | 0/0 | 1/1 | 0/1 | 1/1 | 1/1 |
| Lateral locomotor score | chr1:95,096,740 | C | T | 1/1 | 0/0 | 0/1 | 1/1 | 1/1 | 1/1 | 0/0 | 0/0 | 0/0 | 0/1 | 1/1 | 0/1 | 0/0 | 0/1 | 0/1 | 0/1 | 0/1 | 0/1 | 1/1 | 1/1 |
| Lateral locomotor | chr7:84,269,306 | C | T | 0/0 | 1/1 | 1/1 | 0/0 | 0/0 | 0/0 | 1/1 | 0/1 | 1/1 | 1/1 | 0/0 | 1/1 | 1/1 | 0/0 | 1/1 | 1/1 | 0/0 | 0/0 | 0/0 | 0/0 |

| | | | | | | | | | | | | | | | | | | | | | | | |
|---|---|---|---|---|---|---|---|---|---|---|---|---|---|---|---|---|---|---|---|---|---|---|---|
| score | | T | C | | | | | | | | | | | | | | | | | | | | |
| OF boli | chr18:23,205,496 | T | C | 1/1 | 0/0 | 1/1 | 1/1 | 0/1 | 0/1 | 0/0 | 1/1 | 0/1 | 0/1 | 1/1 | 0/0 | 0/0 | 0/1 | 0/0 | 0/0 | 0/1 | 0/1 | 0/1 | 0/0 |
| OF distance traveled | chr1:135,389,579 | G | A | 0/0 | 1/1 | 0/1 | 0/0 | 0/0 | 0/0 | 0/1 | 1/1 | 1/1 | 1/1 | 0/0 | 1/1 | 0/1 | 0/0 | 1/1 | 1/1 | 0/0 | 0/0 | 0/0 | 0/0 |
| Rearing locomotor score | chr1:107,298,166 | T | C | 1/1 | 0/0 | 0/0 | 1/1 | 1/1 | 1/1 | 0/0 | 0/0 | 0/0 | 0/1 | 1/1 | 0/1 | 0/1 | 1/1 | 0/0 | 0/0 | 1/1 | 1/1 | 1/1 | 1/1 |
| Rearing locomotor score | chr1:95,087,956 | G | A | 1/1 | 0/0 | 0/1 | 1/1 | 1/1 | 1/1 | 0/0 | 0/0 | 0/0 | 0/1 | 1/1 | 0/1 | 0/0 | 0/1 | 0/1 | 0/1 | 0/1 | 0/1 | 1/1 | 1/1 |
| Total locomotor score | chr1:107,298,166 | T | C | 1/1 | 0/0 | 0/0 | 1/1 | 1/1 | 1/1 | 0/0 | 0/0 | 0/0 | 0/1 | 1/1 | 0/1 | 0/1 | 1/1 | 0/0 | 0/0 | 1/1 | 1/1 | 1/1 | 1/1 |
| Total locomotor score | chr1:95,096,740 | C | T | 1/1 | 0/0 | 0/1 | 1/1 | 1/1 | 1/1 | 0/0 | 0/0 | 0/0 | 0/1 | 1/1 | 0/1 | 0/0 | 0/1 | 0/1 | 0/1 | 0/1 | 0/1 | 1/1 | 1/1 |

**Supplemental Table 5.** Relationship between Zhou et al. 2019 and results from the current study.

| topSNP from Zhou et al., 2019 study | topSNP from nearby QTL from current study | LD interval of QTL in current study |
|---|---|---|
| chr1:50,823,354 | Absent | NA |
| chr1:125,692,364 | Absent | NA |
| chr1:165,902,503 | Absent | NA |
| chr2:61,293,430 | Absent | NA |
| chr3:60,419,606 | Absent | NA |
| chr7:79,179,315 | chr7:84,269,306 | 83,274,530-90,251,798 |
| chr18:81,455,034 | Absent | NA |
